# Interferon signaling in the nasal epithelium distinguishes among lethal and common cold respiratory viruses and is critical for viral clearance

**DOI:** 10.1101/2023.12.18.571720

**Authors:** Clayton J. Otter, David M. Renner, Alejandra Fausto, Li Hui Tan, Noam A. Cohen, Susan R. Weiss

## Abstract

All respiratory viruses establish primary infections in the nasal epithelium, where efficient innate immune induction may prevent dissemination to the lower airway and thus minimize pathogenesis. Human coronaviruses (HCoVs) cause a range of pathologies, but the host and viral determinants of disease during common cold versus lethal HCoV infections are poorly understood. We model the initial site of infection using primary nasal epithelial cells cultured at air-liquid interface (ALI). HCoV-229E, HCoV-NL63 and human rhinovirus-16 are common cold-associated viruses that exhibit unique features in this model: early induction of antiviral interferon (IFN) signaling, IFN-mediated viral clearance, and preferential replication at nasal airway temperature (33°C) which confers muted host IFN responses. In contrast, lethal SARS-CoV-2 and MERS-CoV encode antagonist proteins that prevent IFN-mediated clearance in nasal cultures. Our study identifies features shared among common cold-associated viruses, highlighting nasal innate immune responses as predictive of infection outcomes and nasally-directed IFNs as potential therapeutics.

## INTRODUCTION

A major function of the nasal cavity is to induce turbulent airflow of inspired air which mediates deposition and entrapment of debris and infectious particles in mucus before reaching the lung. As a result, all respiratory viruses establish primary infections in the nasal epithelium, which thus serves as a front-line, dynamic defensive barrier with its apical tight junctions and mucociliary clearance function. Virus transmission models suggest that aerosolized viral particles, the mechanism through which most respiratory viruses spread, achieve maximal deposition density in the nasal cavity where viruses initially replicate^1,2^. This is followed by spread to the lower airway, often mediated by aspiration along a nasal/oral-lung aspiration axis, where subsequent lung pathology may occur^3,4^. Alternatively, mucociliary function as well as efficient induction of antiviral innate immune responses in the nasal epithelium may result in local control of viral replication, limited spread to the lower airway and minimal pathogenesis.

The interferon (IFN) signaling pathway is induced at epithelial surfaces following viral recognition resulting in establishment of an antiviral state that restricts viral spread. Respiratory viruses produce double-stranded (ds)RNA as a byproduct of their replication, which is recognized by host sensors such as melanoma differentiation-associated gene 5 (MDA5) and retinoic acid-inducible gene I (RIG-I)^5–7^. These sensors signal via mitochondrial antiviral signaling protein (MAVS) to induce phosphorylation of IFN regulatory factors 3 and 7 (IRF3/7). Activated IRF3/7 translocate into the nucleus where they mediate transcription of type I (IFN-α, IFN-β) and type III (IFN-λ) IFN genes. IFNs are released from infected cells and signal in both autocrine and paracrine fashions via their receptors to induce Janus kinase (JAK) / signal transducer and activator of transcription (STAT) signaling^6,8^. Activated p-STAT proteins translocate into the nucleus where they induce the transcription of hundreds of IFN-stimulated genes (ISGs) with diverse antiviral effector functions which target multiple steps in the viral replication cycle^9–11^. Additional antiviral innate immune pathways important for restriction of viral replication are also induced secondary to dsRNA recognition, including the protein kinase R (PKR) pathway, which results in shutdown of host translation, and the oligoadenylate synthetase (OAS) / ribonuclease L (RNase L) system, which results in cleavage of host and viral RNAs^5^. All three of these dsRNA-induced pathways converge via induction of downstream inflammatory and cell death pathways^12,13^.

Growing literature associates coordinated, early IFN signaling in the nasal epithelium with improved disease outcomes. Indeed, in an influenza infection model designed to replicate natural infection progression from the upper to lower airway, mice lacking functional type III IFN responses were unable to control viral replication in the upper airway and experienced more severe disease^14,15^. Severe acute respiratory syndrome coronavirus 2 (SARS-CoV-2), the causative agent of coronavirus disease 2019 (COVID-19), causes a spectrum of respiratory disease ranging from asymptomatic infections to lethal pneumonia. Sequencing studies of nasopharyngeal swabs acquired from COVID-19 patients have correlated early IFN response signatures in patients’ noses with reduced disease severity, whereas patients with muted ISG profiles tended to have worse outcomes^16–18^. Consistent with antiviral IFN responses in the nose as determinants of disease outcome, nasally delivered IFN has shown promise as an antiviral strategy in various animal models of SARS-CoV-2^19,20^.

It is thus plausible that viruses associated with common cold phenotypes (characterized by self-limited, upper respiratory symptoms such as runny nose, sore throat, nasal congestion, and facial pressure) may replicate predominantly in the upper respiratory tract due to IFN-mediated restriction of spread to the lower airway. Human rhinoviruses (HRVs) are the most common cause of the common cold, associated with 30-50% of annual cases^21^. HRVs replicate robustly and induce IFN in nasal cell infection models. Common cold-associated human coronaviruses (HCoV-229E, -NL63, -OC43, and -HKU1) contribute an additional 15-30% of common cold cases^22^, however, little is known in terms of these viruses’ replication and innate immune induction^23–25^. In contrast to these common cold-associated viruses, the lethal Middle East respiratory syndrome (MERS)-CoV has been primarily associated with lower respiratory tract replication and lethal pneumonia in humans, with a case-fatality rate of 36%^26,27^. Indeed, MERS-CoV employs various strategies to shut down host IFN signaling, including the expression of multiple accessory and nonstructural proteins that directly antagonize host IFN responses^28,29^. SARS-CoV-2 may illustrate an intermediate lethal CoV phenotype, as growing literature has characterized innate immune evasion strategies by SARS-CoV-2 nonstructural and accessory proteins^30–33^. However, the mechanisms behind the observed variability in COVID-19 among individuals remain to be fully characterized. This may be partially determined by IFN responsiveness in the upper airway but is also related to other host factors such as prior exposure and vaccination status. Influenza viruses are yet another class of respiratory viruses that circulate seasonally and cause significant burden on human health (infecting 5-10% of adults each year). Influenza viruses are associated with replication throughout the respiratory tract despite robust IFN induction in most epithelial cell models, and cause typically more severe “flu-like” symptoms which include fever and myalgias^34–37^. Clearly, respiratory viruses differentially interface with host antiviral signaling, and early antiviral innate immune responses in the nasal epithelium may contribute to these variable clinical disease phenotypes during infection.

In addition to innate immune induction, another factor which may be involved in restricting viral replication to the upper airways is temperature. A gradient of increasing temperature is present in the respiratory tract, ranging from 25°C with inspired air at the nares, to 30-35°C as inhaled air is warmed in the nasal cavity and nasopharynx, and finally reaching 37°C or ambient body temperature in the lungs^38,39^. In comparing the replication of common cold-associated HCoVs with lethal SARS-CoV-2 and MERS-CoV in primary nasal epithelial cells, we previously reported that HCoV-229E and HCoV-NL63 replicate more efficiently at 33°C (nasal temperature) than 37°C (lung temperature), while SARS-CoV-2 has an intermediate phenotype, replicating optimally at 33°C only at late time points^40^. MERS-CoV had no temperature preference. This is consistent with HRV studies which have characterized nasal airway temperature as more permissive to replication by the prototypical common cold virus^41,42^. Elevated temperatures have also been suggested to restrict SARS-CoV-2, but not SARS-CoV, in a lower airway infection model^43^.

We utilize a primary cell culture system in which patient-derived nasal epithelial cells are differentiated at an air-liquid interface (ALI) to model infection of the nasal epithelium where respiratory viruses initially replicate. After differentiation, nasal ALI cultures recapitulate various features of the *in vivo* nasal epithelium, including its heterogeneous cellular population and mucociliary clearance function. We infect nasal ALI cultures with a panel of respiratory viruses associated with a spectrum of clinical disease outcomes in humans, including lethal and common cold-associated coronaviruses, human rhinovirus-16 (HRV-16), and a seasonal influenza A isolate, to identify features shared among common cold-associated viruses in this nasal cell model. We compare the temperature-dependent replication kinetics and IFN response patterns of diverse respiratory viruses, as well as the role of IFN signaling and/or viral antagonism of these responses in the control of replication. Our data indicate that temperature-dependent, IFN-mediated restriction of viral replication may be a universal feature of common cold-associated viruses, further emphasizing the role of early innate immunity as a key determinant of disease severity.

## RESULTS

### Replication of common cold-associated viruses is restricted late during infection of primary nasal epithelial cell cultures

After differentiation, pooled-donor nasal ALI cultures were equilibrated at 33°C (nasal airway temperature) and infected at the apical surface with a panel of respiratory viruses: two lethal HCoVs (SARS-CoV-2 and MERS-CoV), two common cold-associated HCoVs (HCoV-229E, HCoV-NL63), human rhinovirus-16 (HRV-16), and a seasonal H1N1 influenza A isolate (IAV/Brisbane/2007, henceforth referred to as IAV). Apical surface liquid (ASL) was collected at 48-hour intervals until 192 hours post infection (hpi) and titrated via plaque assay to characterize viral replication kinetics in nasal cultures. Average shed viral titers for each virus are shown in **Figure 1**. The viral growth curves segregated into two distinct kinetic patterns. Common-cold associated viruses (HCoV-229E, HCoV-NL63, and HRV-16) reached maximal viral titers early (∼48 hpi) followed by rapid reductions in viral titer to nearly the limit of detection (2 log_10_ pfu/mL) by 144 hpi (**Figure 1A**). We thus define viral clearance by nasal epithelial cells as a reduction in viral titer to nearly the limit of detection. In contrast, SARS-CoV-2, MERS-CoV, and IAV did not exhibit this reduction in viral titers (**Figure 1B**). After reaching peak titers (at 48 hpi for MERS-CoV and IAV, and at 144 hpi for SARS-CoV-2), viral titers plateaued with continued apical release of infectious virus. These data suggest that common cold-associated viruses are efficiently cleared by nasal epithelial cells, whereas lethal HCoVs as well as IAV are not. Additionally, the viruses differed in magnitude of replication, with SARS-CoV-2, HCoV-229E, and HRV-16 all replicating to peak titers of ∼6 log_10_ pfu/mL, though SARS-CoV-2 had delayed replication kinetics. HCoV-NL63, MERS-CoV, and IAV reached peak titers 100-fold lower (approximately 4 log_10_ pfu/mL).

**Figure 1.**
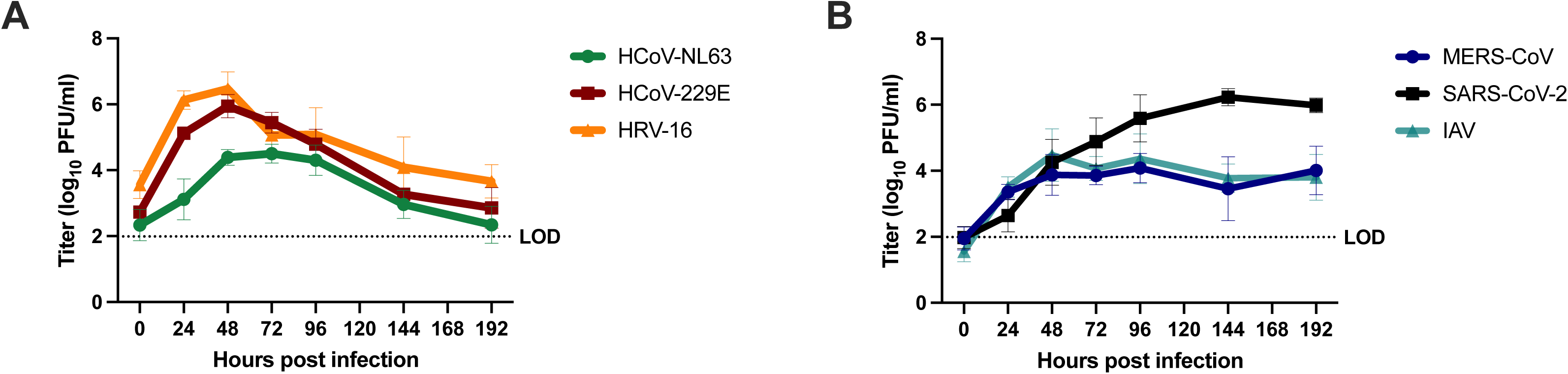
Respiratory viruses exhibit two distinct replication phenotypes in primary nasal epithelial cells. Pooled-donor nasal ALI cultures were infected at the apical surface with the indicated virus (MOI = 1 PFU/cell, 33°C). Apical surface liquid (ASL) was collected at the indicated time points post infection and quantified via plaque assay. Averaged titers from infections in six independent sets of pooled-donor ALI cultures are shown in (A) for HCoV-NL63, HCoV-229E, and HRV-16 and in (B) for SARS-CoV-2, MERS-CoV as mean ± standard deviation (SD). The dotted line indicates the plaque assay limit of detection (LOD).

### Robust IFN signaling responses are induced following infection with common cold-associated viruses as well as IAV

We and others have investigated immune antagonism by lethal HCoVs, particularly by MERS-CoV which adeptly shuts down IFN signaling and other innate immune responses induced following recognition of dsRNA^28,29,44^. In contrast, SARS-CoV-2 activates dsRNA-induced innate immune responses such as IFN signaling and the protein kinase R (PKR) pathway in respiratory epithelial cell lines. However, induction of IFN responses during SARS-CoV-2 infection of primary nasal cells is delayed relative to IAV^37,44^. Thus, we hypothesized that the lack of viral clearance during SARS-CoV-2 and MERS-CoV infection of nasal ALI cultures may be related to impaired innate immune responses relative to those induced following common cold-associated viruses.

We performed bulk RNA sequencing to identify differentially regulated genes (relative to mock-infected cultures) following nasal cell infection by each respiratory virus characterized. Genes with significant up- or down-regulation were assessed using DESeq2 followed by Gene Set Enrichment Analysis (GSEA)^45,46^. This analysis identified immune response genes and specifically the antiviral IFN response as the most significantly upregulated pathway following infection by each of the respiratory viruses except for MERS-CoV (consistent with its efficient shutdown of antiviral responses). Volcano plots highlighting fold changes in normalized transcript abundances relative to mock-infected cultures are shown in **Figure 2A**, with IFN response genes annotated from the Molecular Signatures Database (MSigDB) HALLMARK_INTERFERON_ALPHA_RESPONSE gene list highlighted in green^47^. This analysis confirmed the lack ISG induction by MERS-CoV infection. Additionally, it revealed that relatively few ISGs were induced following SARS-CoV-2 infection compared to the common cold-associated viruses, although more than during MERS-CoV infection. The total number of genes related to the IFN response that reached significance thresholds for each virus is shown in **Figure 2B**. Infection with the common cold-associated HCoVs as well as HRV-16 and IAV induced a larger number of ISGs with greater log_2_ fold change values above mock levels compared to either MERS-CoV or SARS-CoV-2 infection. Though IAV was not cleared by nasal cells, it was associated with robust IFN and ISG induction (in a similar pattern to the common cold-associated viruses), suggesting IAV may exhibit a unique phenotype in this nasal cell model. Stat values (which integrate the magnitude of expression change as well as statistical significance) calculated in DESeq2 for genes in the MSigDB list were combined from each infection and visualized as a heatmap using Morpheus. We performed hierarchical clustering of IFN-related genes to generate a heatmap shown in **Figure 2C**, ranking stat values of each ISG from least (colored blue) to most robustly (colored red) changed during infection with each respiratory virus. Approximately half of the genes involved in IFN signaling were most robustly induced by IAV (with similar induction by HRV-16 for a subset of these). An additional large category of IFN-related genes was strongly induced by all three common cold-associated viruses as well as IAV. In comparison, SARS-CoV-2 and MERS-CoV infected cells demonstrated expression patterns most similar to those of mock-infected cultures.

**Figure 2.**
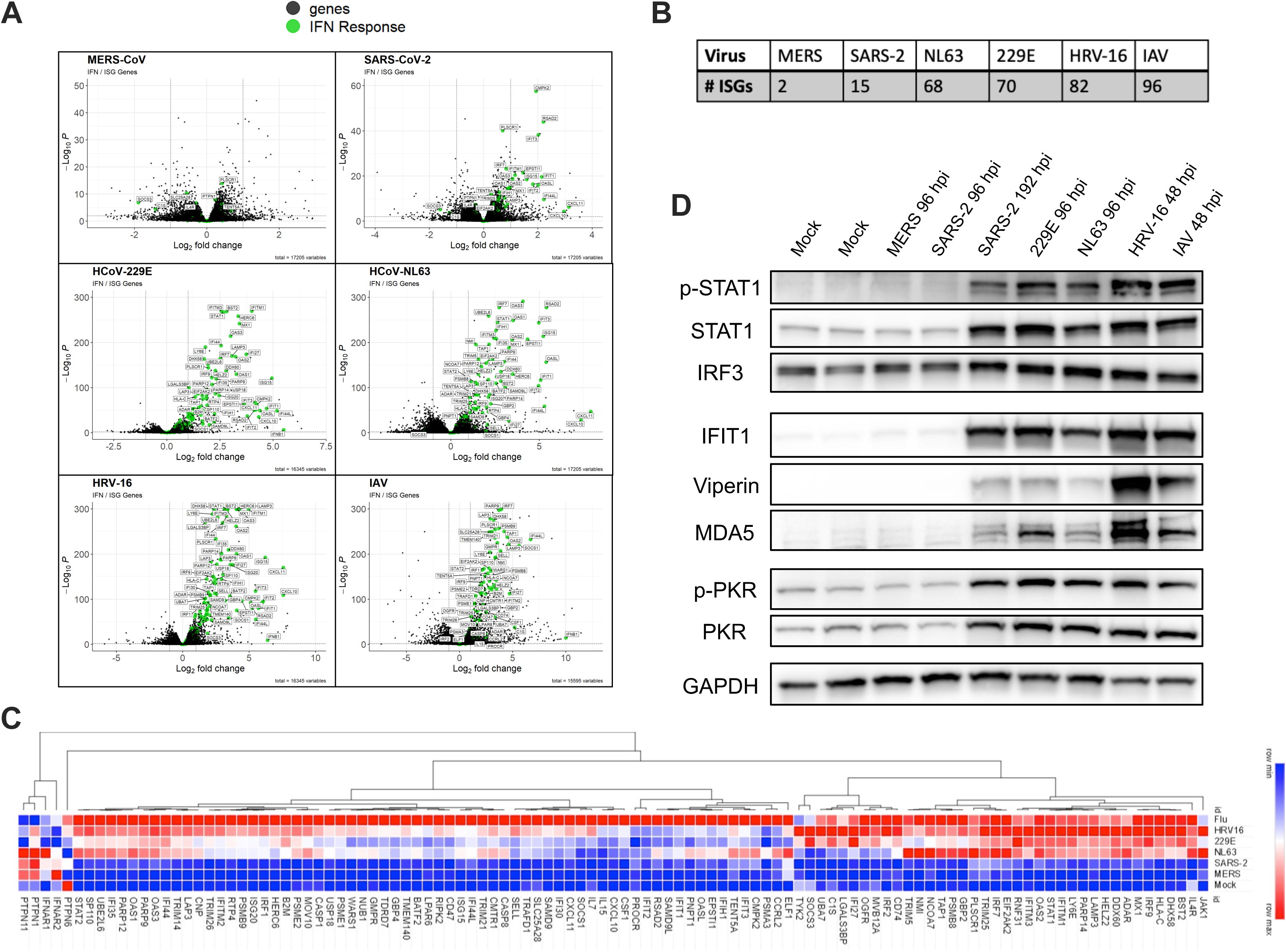
Common cold-associated viruses induce robust, early IFN responses. Nasal ALI cultures were infected with each virus (MOI = 1, 33°C), cells were lysed at 96 hpi (SARS-CoV-2, MERS-CoV, HCoV-229E, HCoV-NL63) or 48 hpi (HRV-16, IAV), RNA extracted and analyzed by RNA-seq. (A) Volcano plots for differentially expressed genes for each virus relative to mock-infected cultures. Genes involved in the IFN signaling response are indicated in green. Significance cutoffs were indicated by dotted lines for both log_2_ fold change values and adjested p-values (padj). (B) The number of ISGs reaching significance thresholds for each respiratory virus was quantified. (C) Heatmap generated via hierarchical clustering of IFN-related genes. Viruses were ranked in terms of degree of induction of each ISG based on DESeq2 stat values, from least upregulated (blue) to most upregulated (red). Data from mock-infected cultures was included and set to row minimum for each gene. (D) Western blot analysis of whole cell lysates collected at indicated times following infection. Time point for this analysis is matched to the time point analyzed via RNAseq, except for SARS-CoV-2, for which an additional sample at 192 hpi was included. Samples were separated via SDS-PAGE followed by transfer on to a PVDF membrane for detection using indicated antibodies.

These RNA-seq patterns were complemented via RT-qPCR quantifying mRNA abundance of type I and III IFN as well as five representative ISGs (**Supplement S1**). To complement the comparison of ISG expression at the RNA level, protein lysates from nasal cells infected with each virus as well as mock-infected cultures were analyzed via western blot to compare levels of STAT1 phosphorylation as well as ISG protein abundances (interferon-induced protein with tetratricopeptide repeats 1, IFIT1, Viperin, MDA5, PKR) among the viruses **(Figure 2D**). For this analysis, early (96 hpi) and late (192 hpi) protein lysates for SARS-CoV-2 were included, given reports of delayed IFN induction by SARS-CoV-2 (RNA-seq analysis for SARS-CoV-2 was at 96 hpi)^37^. Western blots further validate the RNA-seq analysis, highlighting no detectable increase in STAT phosphorylation or ISG protein levels above mock levels following MERS-CoV infection or early during SARS-CoV-2 infection. However, IFN responses do occur late during SARS-CoV-2 infection (192 hpi). Both common cold-associated HCoVs (−229E and -NL63) as well as HRV-16 and IAV resulted in robust STAT phosphorylation as well as robust increases in ISG protein abundances much earlier (48 hpi). We also evaluated activation of another dsRNA-induced antiviral pathway, the PKR pathway and found that patterns of PKR phosphorylation mirrored the gradient of ISG induction. Confirmation of infection via immunoblotting for capsid proteins of each virus is shown in **Supplement S2**. Taken together, common cold-associated viruses, as well as IAV, induce robust IFN signaling responses early during infection of primary nasal cells, whereas MERS-CoV results in no detectable IFN induction and SARS-CoV-2 is associated with delayed IFN responses.

### Inhibition of IFN signaling prevents clearance of common cold-associated viruses, resulting in prolonged viral burden which impacts epithelial barrier integrity

To determine how IFN induction impacts viral replication, we treated nasal cell cultures with ruxolitinib (RUX), a small molecular JAK1/2 inhibitor, which inhibits IFN signaling and induction of ISGs^48^. In vehicle control (DMSO)-treated nasal cells (shown in gray for each virus), common cold-associated viruses (HCoV-229E and -NL63 as well as HRV-16) were efficiently cleared at late time points (**Figure 3A**). However, during infection in the presence of RUX (maroon curves), replication was prolonged and viral clearance did not occur, suggesting that viral clearance is IFN-mediated. In contrast, RUX treatment had only a minor impact on replication of MERS-CoV, consistent with minimal induction of IFN signaling responses by MERS-CoV. SARS-CoV-2 titers were reproducibly increased ∼ten-fold in the presence of RUX only late in infection, consistent with delayed ISG responses by SARS-CoV-2 (**Figure 2D**). Although IAV was associated with robust induction of IFN signaling, RUX treatment had an intermediate effect on viral replication with slightly increased viral titers at late time points (144, 192 hpi).

**Figure 3.**
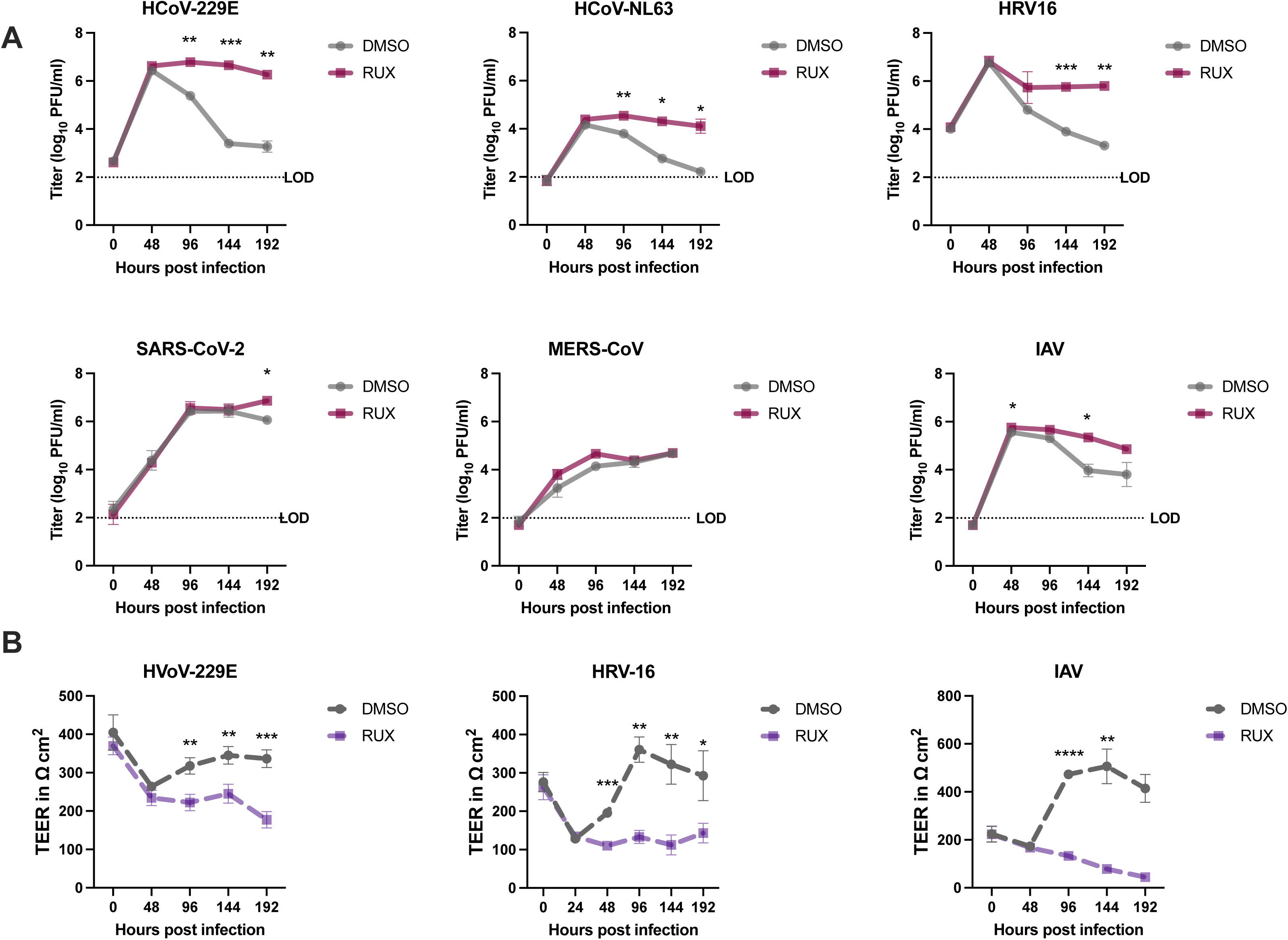
Clearance of common cold-associated viruses is IFN-mediated. Nasal ALI cultures were pre-treated with either ruxolitinib (RUX) or DMSO (vehicle) control at a concentration of 10 µM in the basal media for 48 hours prior to infection, followed by infection in triplicate with the indicated virus (MOI = 1, 33°C). (A) ASL was collected at the indicated time points, released virus quantified via plaque assay, and the average viral titer in each condition is shown as mean ± standard deviation (SD). (B) Trans-epithelial electrical resistance (TEER) was measured prior to infection (0 hpi) and at 48-hour intervals following infection. Average TEER values from triplicate transwells in each condition are shown as mean ± SD. Statistical significance of differences in titer (A) or TEER (B) in RUX-treated vs. control cultures was calculated by repeated measures two-way ANOVA: *, *P* ≤ 0.05; **, *P* ≤ 0.01; ***, *P* ≤ 0.001; ****, *P* ≤ 0.0001. Comparisons that were not statistically significant are not labeled. Data shown is from one experiment representative of three (A) or two (B) independent experiments, each performed in triplicate using pooled-donor nasal ALI cultures derived from four to six individual donors.

Since RUX treatment results in inefficient clearance of common cold-associated viruses (and to some extent IAV), we sought to investigate how this prolonged viral burden impacted epithelial barrier integrity. We previously demonstrated defects in epithelial barrier function as measured by loss of trans-epithelial electrical resistance (TEER) early during HCoV-229E infection in nasal cultures, with recovery of epithelial barrier integrity at late time points, whereas HCoV-NL63 caused TEER defects only at late time points. HRV-16 and IAV have also been associated with impaired epithelial barrier function in nasal cell models^49–51^. Thus, we evaluated TEER at 48-hour intervals in the presence of RUX for HCoV-229E, HRV-16, and IAV-infected nasal cells (**Figure 3B**). For each of these viruses, all infected cultures (vehicle control- or RUX-treated) exhibited early loss of TEER, indicating barrier disruption. While untreated cultures recovered to pre-infection (or higher in the case of IAV) TEER levels, RUX treatment blocked TEER recovery to healthy levels. Thus, prolonged viral burden in the context of impaired antiviral IFN responses (modeled with RUX treatment) has a detrimental impact on the recovery of epithelial barrier integrity.

### Inactivation of IFN antagonists encoded by MERS-CoV and SARS-CoV-2 renders lethal HCoVs to exhibit features of common cold-associated viruses

Since the lethal HCoVs either completely shut down (in the case of MERS-CoV) or significantly delay (in the case of SARS-CoV-2) IFN signaling responses, we hypothesized that inactivation of the IFN antagonists encoded by these viruses would result in IFN-mediated clearance in nasal cell cultures. Both viruses express the conserved CoV nonstructural protein 15 (nsp15), which contains an endoribonuclease activity (EndoU) which limits IFN responses^29,52–54^. MERS-CoV additionally encodes accessory protein NS4a which binds and sequesters dsRNA from recognition from host sensors^55,56^. We have previously reported that mutants of each virus expressing an inactivated EndoU (nsp15^mut^) or a MERS-CoV double mutant additionally lacking expression of NS4a (MERS-CoV-nsp15^mut^/ΔNS4a) induce increased IFN signaling responses^28,29,57^. Western blot analysis of protein lysates from nasal cultures infected with either MERS-CoV-nsp15^mut^/ΔNS4a or SARS-CoV-2-nsp15^mut^ revealed increased IFN signaling responses compared to their respective wild-type (WT) parental viruses, indicated by increased STAT1 phosphorylation as well as increased abundance of representative ISGs IFIT1, viperin, MDA5, and PKR (**Figure 4A, 4B**). The MERS-CoV double mutant resulted in robust induction of the IFN pathway at earlier time points (48 and 96 hpi), whereas the SARS-CoV-2 mutant virus showed IFN signatures most clearly at 192 hpi, consistent with our prior studies^40,57^. After confirming that inactivation of IFN antagonists encoded by the lethal HCoVs resulted in robust IFN induction in nasal cell cultures, we conducted growth curves to determine how increased IFN signaling impacted viral replication. MERS-CoV-nsp15^mut^/ΔNS4a was attenuated for replication compared to WT MERS-CoV beginning at 48 hpi. However, while WT MERS-CoV continued to replicate at late times post infection, MERS-CoV-nsp15^mut^/ΔNS4a titers declined to nearly the limit of detection by 144 and 192 hpi (**Figure 4C**). For the SARS-CoV-2 nsp15 mutant, growth curves were extended to 240 hpi given the delay in IFN induction by SARS-CoV-2 that was still apparent during infection with the mutant virus. SARS-CoV-2 nsp15^mut^ was attenuated relative to WT SARS-CoV-2 only at late time points (beginning at 144 hpi) and similarly exhibited a trend toward clearance by nasal cells at 192 and 240 hpi, while WT SARS-CoV-2 titers did not decline (**Figure 4D**). To confirm that the decline in viral titers for these MERS-CoV and SARS-CoV-2 mutant viruses was IFN-mediated, nasal cells were pre-treated with RUX, resulting in near-complete rescue of viral replication to WT levels for both mutant viruses as we have recently reported for SARS-CoV-2 (**Figure 4C, 4D**)^57^. Thus, inactivation of IFN antagonists encoded by SARS-CoV-2 and MERS-CoV results in a phenotype mirroring common cold-associated viruses, whereby mutant viruses undergo IFN-mediated clearance in nasal cultures.

**Figure 4.**
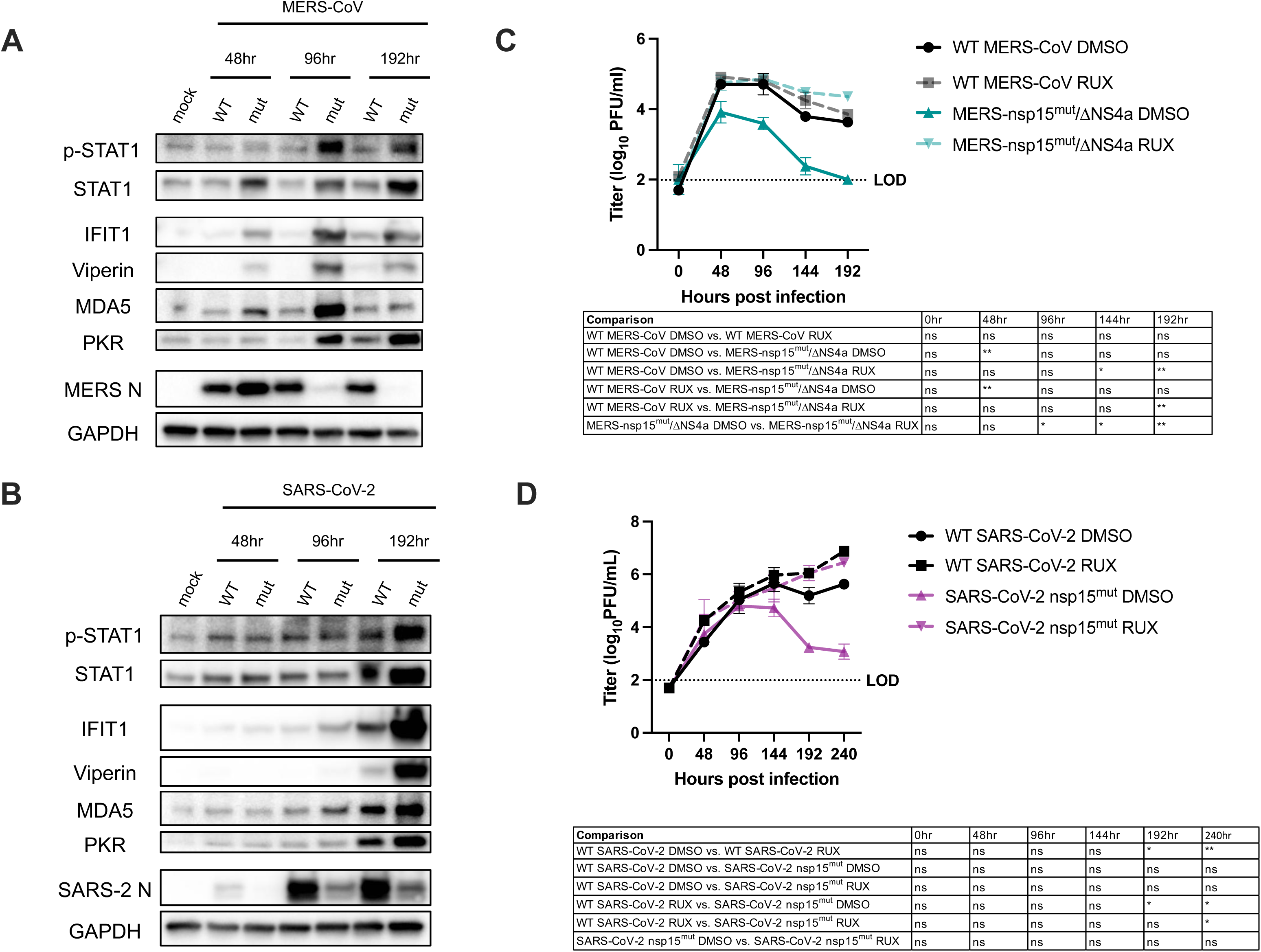
Lethal HCoVs with inactivated IFN antagonists exhibit IFN-mediated clearance. (A-B) Western blot analysis of protein from cells lysed at indicated times post infection with (A) WT MERS-CoV and MERS-nsp15^mut^/ΔNS4a or (B) WT SARS-CoV-2 and SARS-CoV-2 nsp15^mut^. Proteins were separated via SDS-PAGE followed by transfer on to a PVDF membrane for detection using indicated antibodies. (C-D) Nasal ALI cultures were pre-treated with either ruxolitinib (RUX) or DMSO control at a concentration of 10 µM in the basal media for 48 hours prior to infection. Cultures were then infected in triplicate (MOI = 1, 33°C) with either (C) MERS-nsp15^mut^/ΔNS4a or WT MERS-CoV or (D) SARS-CoV-2 nsp15^mut^ and WT SARS-CoV-2. ASL was collected at indicated time points after infection and infectious virus was quantified via plaque assay. Average viral titer for each virus/drug condition is shown as mean ± SD. Statistical significance of differences in titer between each condition was calculated by repeated measures two-way ANOVA and shown as a table: *, *P* ≤ 0.05; **, *P* ≤ 0.01; ***, *P* ≤ 0.001; ****, *P* ≤ 0.0001. Data shown is from one experiment representative of two independent experiments, each performed in triplicate using pooled-donor nasal ALI cultures derived from four to six individual donors.

### Respiratory viruses are differentially sensitive to IFN pre-treatment

Given the association between robust IFN induction and viral clearance, we hypothesized that lethal HCoVs may be particularly sensitive to IFN signaling relative to common cold-associated viruses given that these lethal viruses encode numerous strategies to antagonize IFN responses. Various reports have highlighted a marked sensitivity of SARS-CoV-2 and MERS-CoV to IFN pre-treatment^58,59^, however, few studies have queried the sensitivity of common cold-associated HCoVs to exogenous IFNs. To test this, nasal ALI cultures were pre-treated with type I (IFN-β) or type III (IFN-λ) IFN (100 units/mL in the basal medium) 16 hours prior to infection and then viral replication was quantified at 24-hour intervals until 96 hpi (**Figure 5**). Interestingly, replication of all four HCoVs (SARS-CoV-2, MERS-CoV, HCoV-NL63, and HCoV-229E) was nearly completely inhibited by treatment with either IFN-β or IFN-λ. HRV-16 and IAV, on the other hand, exhibited intermediate sensitivity to IFN pre-treatments. Replication of both HRV-16 and IAV was reduced following IFN-β pre-treatment, though not to the degree observed for HCoVs (in which case, replication was near the limit of detection for all IFN pre-treatments). HRV-16 and IAV were also largely insensitive to IFN-λ, which is consistent with reports that IFN-λ can be less potent than type I IFN in certain contexts^60,61^. These findings may explain the lack of viral clearance during IAV infection despite IAV being associated with the most robust IFN signaling responses in our respiratory virus panel (**Figures 1B, 2D**). Our data highlight the uniform sensitivity of HCoVs to either type I or type III IFN pre-treatment, consistent with IFN-mediated viral clearance for common cold-associated HCoVs and antagonism of IFN responses by the lethal HCoVs.

**Figure 5.**
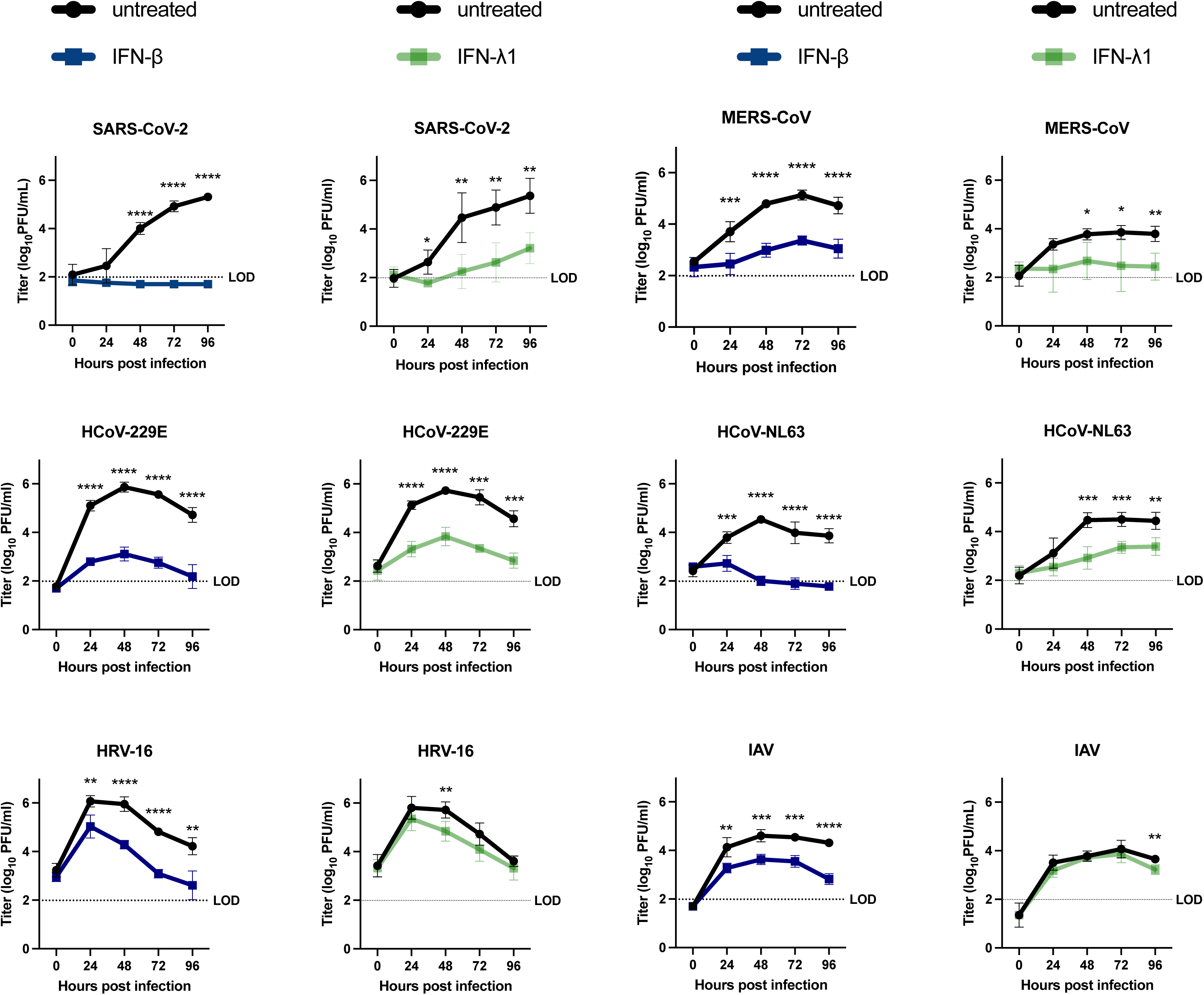
Respiratory viruses are differentially sensitive to IFN pre-treatments in primary nasal epithelial cells. IFN-β or IFN-λ (100 units/ml) was added to the basal media of nasal ALI cultures or control cultures were mock-treated. 16 hours later, cultures were infected with indicated virus (MOI = 1, 33°C). ASL collected at 24-hour intervals following infection was used for quantification of infectious virus release by plaque assay. Average viral titers are shown as mean ± SD, with plaque assay limit of detection (LOD) indicated by dotted line. Statistical significance of differences in average titer in IFN-treated cultures compared to untreated cultures was calculated via repeated measures two-way ANOVA: *, *P* ≤ 0.05; **, *P* ≤ 0.01; ***, *P* ≤ 0.001; ****, *P* ≤ 0.0001. Comparisons that were not statistically significant are not labeled. Data shown is the average of two experiments performed using independent batches of donor nasal ALI cultures, each derived from four to six donors.

### Temperature-mediated defects in replication of common cold-associated viruses are related to IFN signaling

Our prior report comparing HCoV infections in nasal ALI cultures demonstrated that HCoV-229E, HCoV-NL63, and SARS-CoV-2 (at late time points) have a clear preference for replication at 33°C (nasal airway temperature) relative to 37°C (lung temperature), whereas MERS-CoV exhibited no differences in replication if cultures were incubated at either temperature^40^. HRV has also been shown to replicate more efficiently at (33°C), suggesting that this may be another common feature of common cold-associated viruses. The IAV isolate (H1N1 Brisbane/2007) used in this study exhibited no differences in replication if nasal cell infections were conducted at 33°C or 37°C, which was additionally confirmed for MERS-CoV (**Supplement S3**).

Given our data indicating the impact of antiviral IFN responses in restricting viral replication, we sought to determine how IFN responses were regulated by temperature in nasal ALI cultures. We conducted nasal cell infections at 33°C or 37°C with each of the four viruses that exhibited a replication preference for nasal airway temperatures – both common cold-associated HCoVs, HRV-16, as well as SARS-CoV-2 at late time points. Protein lysates from infected cultures were collected at various times post infection for analysis by western blot to compare IFN responses at 33°C with those at 37°C. Representative data is shown for HCoV-NL63 (**Figure 6A**) and SARS-CoV-2 (**Figure 6B**). During HCoV-NL63 infection, levels of STAT1 phosphorylation as well as representative ISGs (IFIT1, Viperin, MDA5) are significantly upregulated when infections are conducted at 37°C relative to 33°C. This increased IFN signaling is most apparent at early time points (24, 48 hpi). Viral replication is relatively similar at 33°C vs. 37°C at these early time points (**Figure 6C**), however, enhanced IFN responses at 37°C mediate more efficient restriction of viral replication (relative to 33°C), resulting in more rapid viral clearance. An observable “switch” then occurs, whereby prolonged viral replication at 33°C results in relatively higher IFN responses compared to 37°C at later time points (96, 144 hpi), after IFN-mediated clearance occurs at 37°C. For SARS-CoV-2, the most robust STAT1 phosphorylation as well as downstream ISG protein expression occurred at 144 hpi at 37°C, the time point immediately prior to the observed growth defect of SARS-CoV-2 at 37°C **(Figure 6D**). Evidence of increased IFN response following SARS-CoV-2 infection at 37°C relative to 33°C is apparent as early as 96 hpi. When SARS-CoV-2- infected nasal cells were incubated at 33°C, IFN signaling responses never reached the level of induction observed during infections conducted at 37°C. Similar data for HCoV-229E and HRV-16 are shown in **Supplement S4**, illustrating enhanced IFN responses at 37°C during infection with each of these common cold-associated viruses.

**Figure 6.**
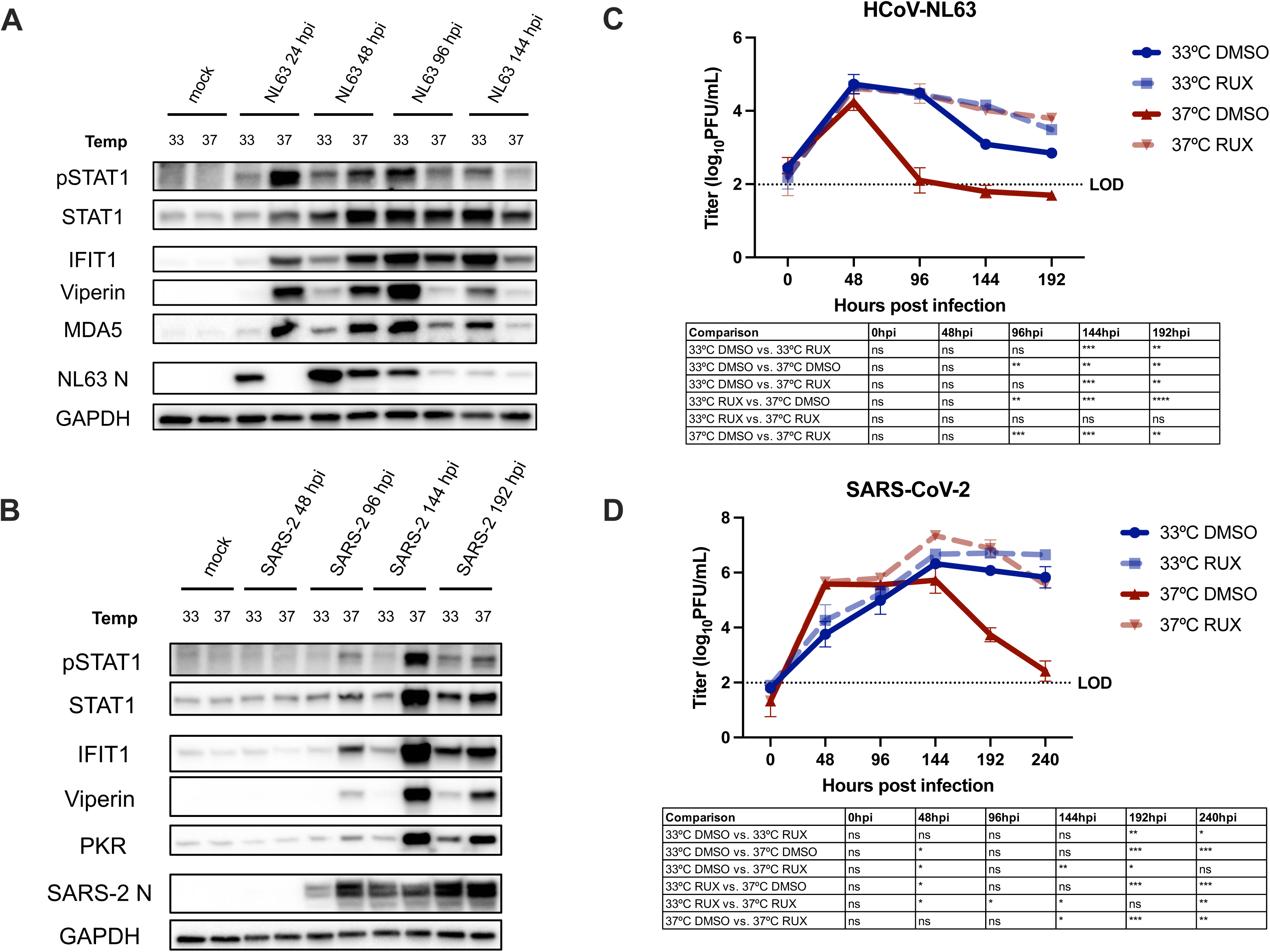
Enhanced IFN responses restrict replication of common cold-associated viruses at elevated temperature. (A,B) Nasal ALI cultures were equilibrated at indicated temperature (33°C or 37°C) for 48 hours prior to infection by HCoV-NL63 (A) or SARS-CoV-2 (B). Western blot analysis was performed using lysates of cells harvested at indicated time points. Immunoblots were probed with antibodies against indicated proteins involved in the IFN signaling response. Data shown are from one experiment representative of three independent experiments conducted in separate batches of pooled-donor nasal cultures. (C,D) Cultures were pre-treated with either RUX or DMSO at 10 µM in the basal media at the start of temperature equilibration 48 hours pre-infection. Cultures were then infected with HCoV-NL63 (C) or SARS-CoV-2 (D) in triplicate (MOI = 1), ASL collected at indicated time points and infectious virus quantified by plaque assay. Average viral titer for is shown as mean ± SD. Statistical significance of differences in titer between each condition was calculated by repeated measures two-way ANOVA and shown as a table: *, *P* ≤ 0.05; **, *P* ≤ 0.01; ***, *P* ≤ 0.001; ****, *P* ≤ 0.0001. Data shown are from one experiment representative of three independent experiments, each performed in triplicate using independent batches of pooled-donor cultures.

To determine if temperature-dependent IFN responses contributed to restriction of viral replication at 37°C, we further conducted nasal cell infections at each temperature in the presence or absence of RUX to impair IFN signaling (**Figure 6B, 6D**). The defect in replication during HCoV-NL63 infection at 37°C is completely rescued to the levels of virus observed at 33°C in cells that were treated with RUX. Indeed, viral titers in the 33°C RUX and the 37°C RUX treated conditions were not significantly different at any time during HCoV-NL63 infection (**Figure 6B**). RUX treatment similarly rescued the temperature-mediated growth defect of HCoV-229E (**Supplement S4C**). Given the delayed IFN induction by SARS-CoV-2, as well as its delayed temperature phenotype (exhibiting preferred replication at 33°C only at very late time points concurrent with IFN induction), growth curves for SARS-CoV-2 were extended to 240 hpi (**Figure 6D**). Although RUX treatment had minimal impact on SARS-CoV-2 titers at 33°C (consistent with data shown in **Figure 4**), it had a more dramatic impact on SARS-CoV-2 replication at 37°C. Similar to observations for HCoV-NL63 and HCoV-229E, RUX treatment nearly completely rescued the defect in replication at 37°C during SARS-CoV-2 infections at late time points. Interestingly, RUX treatment did not similarly rescue the temperature-mediated growth defect of HRV-16, though a slight increase in viral titer was observed when HRV-16-infected cultures were incubated at 37°C in the presence of RUX (relative to 37°C vehicle control-treated cultures) (**Supplement S4D**). RUX treatment had a much more robust impact on HRV-16 titers at 33°C,resulting in complete impairment of viral clearance (**Figure 3A**). Taken together, heightened IFN signaling responses at 37°C mediate restriction of common cold-associated HCoVs as well as SARS-CoV-2 at late time points in primary nasal ALI cultures and thus contribute to optimal replication of these viruses at nasal airway temperatures.

### The omicron BA.1 variant of SARS-CoV-2 exhibits some, but not all, features of common-cold associated viruses

The dominant SARS-CoV-2 omicron variant of concern (VOC) has been associated with heightened replication in the upper respiratory tract, increased transmissibility, as well as a propensity to cause common cold-like symptoms (such as runny nose and sore throat)^62,63^. Thus, we compared the ancestral SARS-CoV-2/WA-1 (used for all SARS-CoV-2 experiments previously described) with omicron BA.1 in this nasal cell model to determine if SARS-CoV-2 had evolved to exhibit features of common cold-associated viruses. Comparing the replication kinetics of SARS-CoV-2 WA-1 with omicron BA.1 at 33°C confirmed a number of reports which have found that omicron replicates more rapidly in nasal cultures^64,65^. Omicron BA.1 replication reached peak titers by 48 hpi, whereas SARS-CoV-2 WA-1 titers peaked at 144 hpi (**Figure 7A**). Though we initially hypothesized that omicron BA.1 would exhibit increased temperature sensitivity with a more significant decline in viral titers than occurs during SARS-CoV-2 WA-1 infection at 37°C, we found in contrast that omicron BA.1 replication was markedly insensitive to temperature when comparing replication at 33°C vs 37°C (**Figure 7A**). Omicron titers remained at peak levels at late time points (as late as 240 hpi) when infections were conducted at either temperature, suggesting that omicron BA.1 does not undergo clearance by nasal epithelial cells as we observed for common cold-associated viruses.

**Figure 7.**
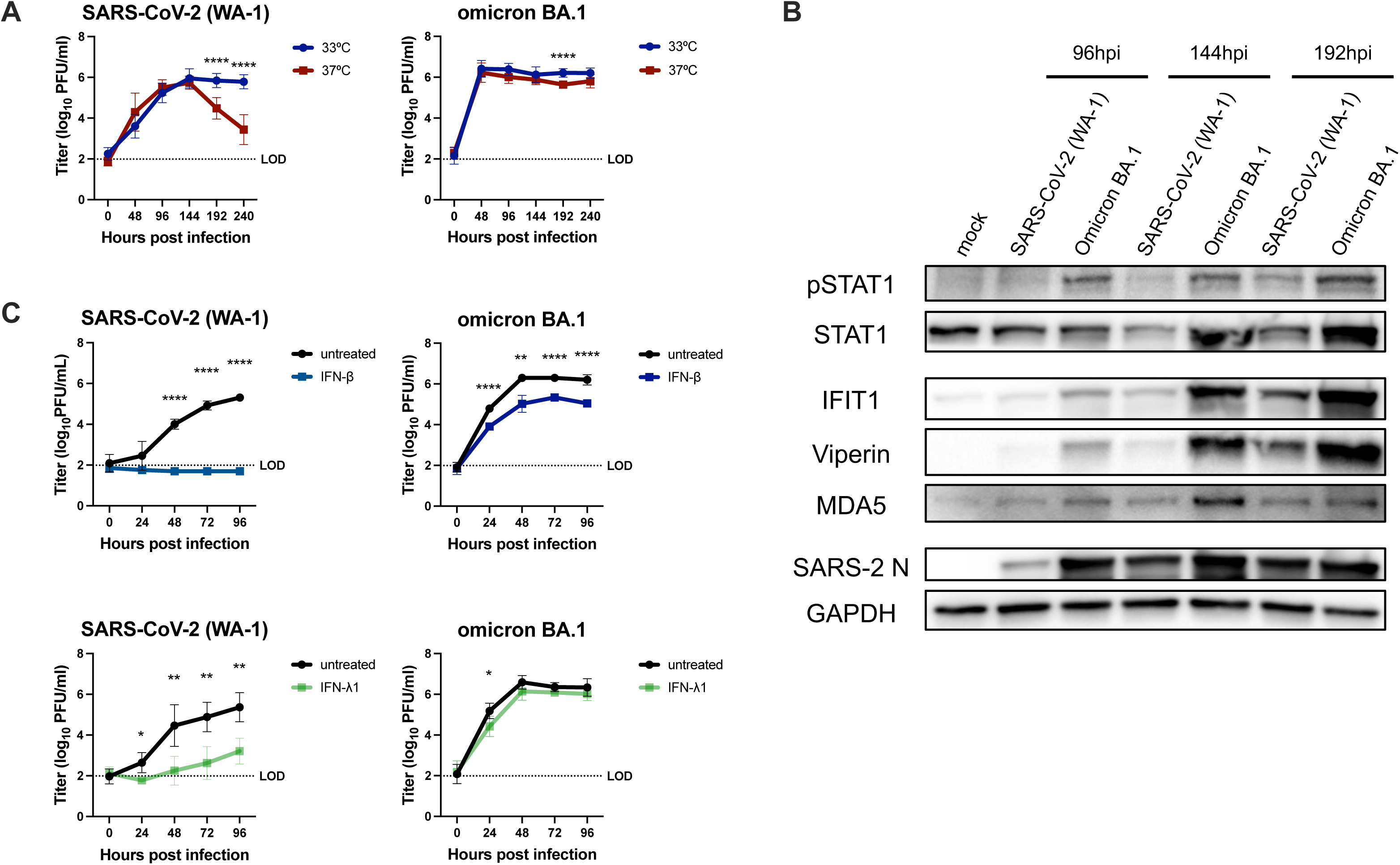
Omicron BA.1 exhibits a unique phenotype in primary nasal epithelial cells. (A) Nasal ALI cultures were equilibrated at the indicated temperature for 48 hours and then infected with either SARS-CoV-2 (WA-01) or omicron BA.1 (MOI = 1). ASL was collected at indicated times and released infectious virus quantified by plaque assay. Data shown are the average from three independent experiments. (B) Western blot analysis was performed using whole cell lysates collected at indicated time points after infections with SARS-CoV-2 WA-01 or omicron BA.1 (MOI = 1, 33°C). Data shown are from one representative of four experiments conducted in independent batches of pooled-donor nasal cultures. (C) Nasal ALI cultures, pre-treated with IFN-β or IFN-λ for 16 hours were infected with either SARS-CoV-2 (WA-01) or omicron BA.1 (MOI = 1, 33°C), ASL was collected at 24-hour intervals and quantified for infectious virus by plaque assay. Data shown is the average of two independent experiments. Statistical significance of differences in titer for each virus at 33°C vs. 37°C (A) or in IFN-treated vs. untreated cultures (C) was calculated via repeated measures two-way ANOVA : *, *P* ≤ 0.05; **, *P* ≤ 0.01; ***, *P* ≤ 0.001; ****, *P* ≤ 0.0001. Comparisons that were not statistically significant are not labeled.

To assess the degree of IFN signaling pathway induction during omicron infection, we performed western blot analysis using protein extracted from SARS-CoV-2 WA-1- or omicron BA.1-infected nasal cells, which revealed an increase in STAT1 phosphorylation as well as ISG protein abundances during omicron BA.1 infection relative to SARS-CoV-2 WA-1 (**Figure 7B**)^65^. Finally, since clearance of common cold-associated viruses was associated with IFN responses, we compared the sensitivity of omicron BA.1 with the earlier isolate, SARS-CoV-2 WA-1 (**Figure 7C**). Pre-treatment with either type I (IFN-β) or type III (IFN-λ) IFN was strongly antiviral against SARS-CoV-2 WA-1, nearly completely restricting viral replication. However, omicron BA.1 was much less sensitive to either of these IFN pre-treatments, replicating to high titers in the presence of strong and sustained IFN responses in primary nasal cells. Thus, the omicron BA.1 variant has a unique phenotype, whereby it induces stronger IFN responses in nasal cell cultures than SARS-CoV-2 WA-1 but exhibits reduced IFN sensitivity and therefore does not illustrate temperature sensitivity nor does it undergo IFN-mediated clearance.

## 5.4 DISCUSSION

Efficient induction of antiviral IFN signaling in the nasal epithelium has the potential to restrict the spread of respiratory viruses to the lower airway, thus preventing the development of more severe lung disease. Viruses such as HRV-16, HCoV-229E, and HCoV-NL63 are typically associated with upper respiratory tract replication and symptoms, as well as early resolution of infection. Reports of IFN-mediated restriction of HRVs, as well as recent nasopharyngeal sequencing studies of COVID-19 patients that correlate early nasal IFN induction with reduced disease severity suggest that IFN responses in the nasal epithelium may be critical determinants of disease course^16–18,66,67^. Relatively few studies have utilized nasal cell models to compare common cold-associated viruses with more lethal respiratory viruses. Additionally, the common cold-associated HCoVs have been largely overlooked in respiratory virus research due to difficulties using these viruses in traditional cell culture systems as well as minimal interest prior to the COVID-19 pandemic. Thus, we have compared a panel of respiratory viruses spanning three virus families and encompassing diverse clinical phenotypes during human infection in a primary nasal ALI culture model. HRV-16, a prototypical common cold virus of the *Picornaviridae* family, as well as the alphacoronaviruses HCoV-229E and HCoV-NL63, are used to model common cold-associated infections, while lethal betacoronaviruses SARS-CoV-2 and MERS-CoV, as well as the *Orthomyxoviridae* family member IAV (H1N1/Brisbane 2007) serve as comparators associated with more severe respiratory disease. We identified and characterized three features of common cold-associated viral infections in our nasal cell model: robust, early induction of IFN responses, IFN-mediated clearance, and restriction of viral replication at elevated temperatures.

The replication cycle of all three common cold-associated viruses was characterized by an early peak in viral replication followed by clearance to the limit of detection at late time points post infection (**Figure 1A**). Abrogation of IFN signaling with RUX rendered nasal cells unable to clear common cold-associated viruses, suggesting that restriction of viral replication is IFN-mediated (**Figure 3**). The nasal ALI cultures are composed of various epithelial cell populations (ciliated, goblet, and basal cells), but notably do not contain resident innate immune cell populations such as macrophages or dendritic cells that would be present in an *in vivo* nasal epithelium. This suggests that efficient induction of antiviral IFN responses by nasal epithelial cells may be sufficient to significantly limit replication of these viruses and curtail further viral spread, which likely contributes to reduced disease severity. It is also plausible that early IFN responses by nasal epithelial cells suppress initial viral replication, allowing for the recruitment of myeloid cell populations to the nasal epithelium which additionally contribute to viral clearance.

In contrast to this IFN-mediated clearance following infection with common cold-associated viruses, nasal cultures infected with SARS-CoV-2, MERS-CoV, or IAV were unable to clear these viruses (**Figure 1B**). This likely results at least in part from shutdown (for MERS-CoV) or significant delay (for SARS-CoV-2) of IFN responses. Indeed, infections with SARS-CoV-2 or MERS-CoV mutants with inactivated or deleted IFN antagonists resulted in a phenotype similar to that observed for HRV-16 and the common cold-associated HCoVs – with robust IFN/ISG responses that significantly restricted viral replication (**Figure 4**). We hypothesize that evasion of antiviral IFN responses is a crucial factor that allows these lethal HCoVs to replicate in the nasal epithelium and subsequently spread to the lower airway. An interesting parallel can be drawn for MERS-CoV, which causes predominantly upper respiratory tract illness in its animal reservoir, dromedary camels^68,69^. Induction of antiviral innate immunity in the camel upper airway has yet to be characterized, and this may be responsible for the restriction of MERS-CoV replication in camels to the upper airway, resulting in a common cold-like phenotype and limited spread to the camel airway. Receptor distribution may also contribute to this phenomenon, as comparative analysis of the MERS-CoV receptor, dipeptidyl peptidase 4 (DPP4), in camel and human airways has shown high expression of DPP4 in the camel upper airway^70,71^.

Exemplifying an intermediate phenotype, SARS-CoV-2 infection induces delayed IFN responses in the nasal cell model (**Figure 2D**), with pronounced IFN induction similar to that seen early during common cold-associated virus infections, although not detected until 192 hpi. As a result of this delayed IFN induction, SARS-CoV-2 replication is sustained at late times post infection. Our data are consistent with a prior study that described IFN induction by SARS-CoV-2 as delayed relative to IAV in nasal cells^37^. Interestingly, when we conducted SARS-CoV-2 infections at 37°C, we observed earlier and more robust ISG responses and a subsequent trend toward viral clearance (**Figure 6**). The timing of IFN responses has been shown to be a critical determinant of pathogenesis, whereby IFN induction too late following infection can worsen outcomes in a mouse model of MERS-CoV infection^72,73^. Thus, characterizing this delay in IFN induction during SARS-CoV-2 infection may provide insight into the marked variability in COVID-19 severity. The mechanism(s) contributing to delayed IFN induction by SARS-CoV-2 have not been fully elucidated but is likely at least partially due to its various IFN antagonists, including the conserved CoV nsp15 EndoU. When we infect with a SARS-CoV-2 mutant lacking nsp15 EndoU activity, SARS-CoV-2 undergoes IFN-mediated clearance (**Figure 4D**) ^57^. In addition, accessory proteins encoded in genes ORF6 and ORF8 have been reported to have IFN antagonist functions^30,32,74^. For example, the SARS-CoV-2 ORF6 protein inhibits IFN signaling via blockade of STAT protein translocation into the nucleus, a process that is important for induction of IFN transcription ^31,74–78^. Thus, our data indicate that modulating IFN responses via changes in temperature or via inactivation of SARS-CoV-2 IFN antagonists resulted in SARS-CoV-2 exhibiting features of common cold-associated viral infections in nasal epithelial cells.

In addition to viral immune antagonism, our data substantiate a role for temperature-dependent regulation of IFN responses as another factor that may contribute to restriction of common cold-associated viruses to the upper airway. In vitro transcription models posit that small but physiologic increases in temperature can impact alternative splicing and other signaling events leading to increased IFN pathway induction at higher temperatures^79^. HRV-16 as well as both common cold-associated HCoVs included in this study replicated more efficiently at 33°C, exhibiting enhanced relative clearance at 37°C. This is consistent with a study using a mouse-adapted rhinovirus that suggests enhanced IFN responses at 37°C (lung temperature) compared to 33°C (nasal airway temperature) limit rhinovirus replication^41,42^. SARS-CoV-2 also had a preference for replication at 33°C at late times, concurrent with its delayed IFN signature. Interestingly, while RUX treatment to inhibit IFN signaling led to nearly complete rescue of this temperature-dependent attenuation for SARS-CoV-2, HCoV-NL63, and HCoV-229E, only partial restoration of replicaton was observed for HRV-16 (**Supplement S4**). This observation suggests that other factors may contribute to the replication defect of HRV-16 at 37°C. This could be related to cell death pathways or other dsRNA-induced antiviral pathways, such as RNase L and PKR which have been associated with temperature during HRV infections^42^. However, OASs as well as PKR, the dsRNA sensors responsible for RNase L and PKR pathway activation, respectively, are ISGs upregulated following IFN induction, and thus these pathways are likely minimally activated in the presence of RUX. Thus, it is likely that additional virus-related factors related to virion stability as well as virus-encoded enzyme function (viral polymerases or proteases, for example) may also contribute to temperature-mediated replication differences in addition to IFN signaling. *In vitro* studies of the IAV RNA-dependent RNA polymerase function propose that temperature can regulate viral replication machinery, though the impact of temperature on viral enzyme function has not been investigated during authentic infection^80,81^. Although additional mechanisms likely contribute, IFN responses that vary along physiologic airway temperatures may be critical in restricting common cold-associated viruses from spreading to the lower airway, especially during HCoV infection in which RUX treatment rescued temperature defects in our model. Future studies will compare temperature-dependent IFN-mediated restriction of common cold-associated viruses using epithelial cells derived from the upper and lower airway to determine if this mechanism is more robust in nasal cells.

The seasonal IAV isolate used in this study (H1N1 Brisbane/2007) illustrates a unique phenotype. It resembles common cold-associated viruses with robust, early IFN induction, however, these IFN responses were insufficient to mediate viral clearance, and IAV also did not preferentially replicate at nasal airway temperatures. This may be due to decreased sensitivity to IFN responses (especially IFN-λ). Strong induction of IFN occurred, surprisingly, despite IAVs encoding a potent and well-characterized IFN antagonist, the NS1 protein, which possesses dsRNA-binding function and also limits IFN signaling induction^82–85^. Future studies with an IAV mutant lacking expression of this NS1 protein would be informative, as we hypothesize that such a virus may illustrate IFN-mediated temperature sensitivity as well as clearance by nasal epithelial cells, mirroring phenotypes for SARS-CoV-2 and MERS-CoV mutants lacking their IFN antagonists. Analysis of more recent omicron subvariants in our nasal model may indicate further evolution of SARS-CoV-2 with respect to its interaction with the IFN signaling response.

Sensitivity to IFN pre-treatments was a universal feature among all four HCoVs used in this study. However, reduced IFN sensitivity was also observed for the SARS-CoV-2 omicron BA.1 VOC in our model. Though most studies have emphasized mutations in the spike proteins of the SARS-CoV-2 variants, which mediate antibody escape and increased transmissibility, a growing number of reports identify differences in innate immune induction and IFN sensitivity among VOCs. Indeed, the still-dominant omicron variant replicates more rapidly in upper airway models and seems to replicate less efficiently than early SARS-CoV-2 isolates in lung-derived epithelial cell lines such as Calu3 and lower airway ALI cultures^64,86,87^. Though the degree of IFN induction during omicron infection relative to early SARS-CoV-2 isolates is a topic of debate^88,89^, a consensus has emerged in the literature that suggests SARS-CoV-2 VOCs (and especially omicron BA.1) have enhanced IFN resistance, which is consistent with our data^87,90–92^. This reduced IFN sensitivity likely explains the lack of clearance and temperature sensitivity observed during omicron BA.1 infection in our nasal cell model, which parallels our observations during IAV infection. Mutations associated with IFN resistance in SARS-CoV-2 VOCs have yet to be clearly identified, although mutations in nonstructural proteins nsp3, nsp6, and nsp12 as well as accessory gene ORF6 may play a role^31,92^. Such differences in IFN sensitivity represent another factor that likely contributes to infection outcome and early restriction of viral replication in the nasal epithelium.

Leveraging a primary nasal cell model to compare respiratory viruses associated with a range of clinical pathologies, we propose that common cold-associated viruses induce temperature-regulated IFN responses that restrict viral replication. IFN antagonism by lethal HCoVs and reduced IFN sensitivity in the case of IAV and the omicron BA.1 variant of SARS-CoV-2 hinder viral clearance in nasal cell cultures. It would be informative to evaluate this model with additional respiratory viruses, both common cold-associated viruses (such as the paramyxovirus respiratory syncytial virus and respiratory adenoviruses) as well as additional lethal viruses (pandemic strains of IAV as well as SARS-CoV). Early comparisons of the IFN sensitivity of SARS-CoV and SARS-CoV-2 suggest that SARS-CoV-2 was more IFN-sensitive^58^, so we hypothesize that SARS-CoV would not undergo IFN-mediated clearance in nasal cells, irrespective of the degree of IFN induced. This would correlate with heightened severity of clinical disease in SARS-CoV infection relative to SARS-CoV-2. A model whereby highly lethal HCoVs such as MERS-CoV and SARS-CoV either shutdown IFN responses or are insensitive to IFN responses in the nasal epithelium may allow for uninhibited dissemination to the lower airway where they cause more severe respiratory disease.

Thus, antiviral IFN responses in the nasal epithelium mediate early control of viral replication and limit spread to the lower airway and are likely key determinants of respiratory viral disease. Our comparative approach characterizing innate immune responses during diverse respiratory virus infections improves our understanding of the mechanisms differentiating common cold-associated viral infections from those associated with more severe disease. Awareness of these virus-host interactions will help to define risk factors associated with severe disease and allow for the development of antiviral therapies that decrease the global burden of respiratory viruses.

## Acknowledgments

We thank members of the Weiss lab for feedback and discussion of this project. We thank Dr. Anthony Fehr and Dr. Luis Martinez-Sobrido for construction of MERS-CoV and SARS-CoV-2 recombinant viruses, Dr. Scott Hensley for providing the seasonal influenza isolate as well as MDCK cells used in the study, Dr. Andrew Vaughan for providing an additional source of MDCK cells. We thank Drs. David W. Kennedy, James N. Palmer, Nithin D. Adappa, and Michael A. Kohanski for aid in the collection of nasal tissue for establishing primary nasal epithelial cultures. This work was supported by National Institutes of Health grants R01 AI140442 (SRW), R01AI169537 (SRW&NAC), Department of Veterans Affairs Merit Review 1-I01-BX005432-01 (NAC&SRW), the Penn Center for Research on Coronaviruses and Other Emerging Pathogens (SRW). CO was supported in part by F30AI172101 and T32AI055400 and AF in part by T32AI007324.

## Respective Contributions

Designed research: CJO, NAC, SRW

Performed research: CJO, DMR, LHT

Contributed new reagents/analytic tools: DMR, LHT

Analyzed data: CJO, DMR, AF, NAC, SRW

Wrote manuscript: CJO

Revised manuscript: CJO, AF, DMR, NAC, SRW

## MATERIALS AND METHODS

### Growth and differentiation of nasal air-liquid interface (ALI) cultures

Primary nasal cells were obtained via nasal cytologic brushing of patients in the Department of Otorhinolaryngology-Head and Neck Surgery, Division of Rhinology at the University of Pennsylvania and the Philadelphia Veteran Affairs Medical Center after obtaining informed consent. The full study protocol, including the acquisition and use of nasal specimens, was approved by the University of Pennsylvania Institutional Review Board (protocol #800614) and the Philadelphia VA Institutional Review Board (protocol #00781). Any patient with a history of systemic disease or who had recently taken immunosuppressive medications was excluded. After specimen acquisition, nasal ALI cultures were grown and differentiated on semipermeable transwell supports containing 0.4 μm pores as previously described^40,93–95^. Pooled-donor nasal cultures were used for all infections in this study. Nasal epithelial cells derived from four or six individual donors were mixed in equal combinations prior to seeding on to transwell supports. Nasal cells were then grown to confluence with Pneumacult-Ex Plus growth medium present both apically and basally. After reaching confluence, apical medium was removed, and Pneumacult-ALI medium was used in the basal compartment to differentiate nasal ALI cultures. Basal media was replaced two times per week throughout the differentiation period (4 weeks total). All reagents used for nasal ALI culture growth and differentiation were acquired from Stemcell Technologies. Nasal ALI cultures were grown and differentiated at 37°C, followed by equilibration at either 33°C or 37°C for 48 hours prior to infection (indicated in figure legends).

### Viruses

SARS-CoV-2 (USA-WA1/2020 strain) obtained via the Biodefense and Emerging Infections Research Resources Repository (BEI) was propagated in Vero-E6 cells. MERS-CoV was derived from a bacterial artificial chromosome (BAC) encoding the full-length MERS-CoV genome (HCoV-EMC/2012) and was propagated in Vero-CCL81 cells. HCoV-NL63 was propagated in LLC-MK2 cells. HCoV-NL63 stock underwent ultracentrifugation through a 20% sucrose layer to concentrate virus stock for infections as previously described^96^. HCoV-229E (ATCC-VR-740) was propagated in Huh7 cells. HRV-16 (ATCC-VR283) was propagated in HeLa cells. IAV (H1N1, Brisbane/2007) was a kind gift from the laboratory of Dr. Scott Hensley and was propagated on MDCK cells. SARS-CoV-2 and MERS-CoV recombinant viruses were generated SARS-CoV-2 and MERS-CoV BACs as previously described^29,57,97^. All virus stocks were sequenced and compared to wild-type reference sequences available via NCBI. All virus manipulations and infections that involved SARS-CoV-2 and MERS-CoV were conducted in a biosafety level 3 (BSL-3) facility following appropriate and approved personal protective equipment and protocols.

### Infections and quantification of apically shed virus

Viruses were diluted in serum-free Dulbecco’s modified Eagle’s medium (DMEM) to achieve MOI = 1 PFU/cell in a total inoculum volume of 50 μl. For IAV infections, trypsin TPCK at a concentration of 2 μg/ml was added to the inoculum to mediate initiation of infection. Viral inocula were added to the apical compartment of each transwell and allowed to adsorb for 1 hour with rocking. After 1 hour, cells were washed three times with phosphate-buffered saline (PBS), and a fourth PBS wash was collected to confirm adequate removal of input virus (0 hpi time point). At indicated times following infection, airway surface liquid (ASL) was collected via addition of 200 μl of PBS to the apical compartment. ASL samples were subsequently used for quantification of apically shed infectious virus via standard plaque assay as previously described^40,44,95,96^. A different cell line, incubation period, and temperature was used for titration of each virus, as described in **Table 1**.

**Table 1.** Plaque assay conditions for each respiratory virus.

| Virus | Cell line used for infectious virus quantification | Temperature of plaque assay | Incubation period |
| --- | --- | --- | --- |
| SARS-CoV-2 | VeroE6 | 37°C | 3 days |
| MERS-CoV | VeroCCL81 | 37°C | 3 days |
| HCoV-229E | Huh7 | 33°C | 3 days |
| HCoV-NL63 | LLC-MK2 | 33°C | 6 days |
| HRV-16 | HeLa | 33°C | 5 days |
| IAV | MDCK | 37°C | 3 days |

### Cell lines used for quantification of infectious virus by plaque assay

VeroE6 cells (ATCC), VeroCCL81 cells (ATCC), HeLa cells (ATCC) and MDCK cells (gifted from Dr. Andy Vaughan and Dr. Scott Hensley) were cultured in Dulbecco’s modified Eagle’s medium (DMEM) containing 10% L-glutamine and 4.5g/L D-glucose (Gibco, ThermoFisher) supplemented with 10% heat inactivated fetal bovine serum (FBS) (Hyclone, Cytiva) and 1X penicillin/streptomycin (pen/strep) (Gibco, ThermoFisher). Huh7 cells (ATCC) were grown in the same media supplemented additionally with 1X non-essential amino acids (Gibco). LLC-MK2 cells were cultured in minimal essential media (MEM)-α supplemented with 10% FBS.

### Bulk RNA sequencing and analysis

Nasal cultures were infected at MOI = 1 with the indicated viruses. Cells were lysed at 96 hpi (MERS-CoV, SARS-CoV-2, HCoV-229E, HCoV-NL63) or 48hpi (IAV and HRV-16) using RLT Plus buffer and total RNA was extracted using Qiagen RNeasy Plus Mini kit (cat. No. 74004). Samples were sequenced by Azenta Life Sciences with Illumina HiSeq PE 2x150. Read quality was assessed using FastQC v0.11.2^98^. Raw sequencing reads from each sample were quality and adapter trimmed using BBDuk 38.73^99^. The reads were then mapped to the human genome (hg38 with Ensembl v98 annotation) using Salmon v0.13.1^100^. Differential expression between mock and infected experimental conditions were analyzed using the raw gene counts files by DESeq2 v1.22.1^45^. Significance thresholds for significantly altered gene expression were set a log_2_ fold change value of > 1 or < -1 and a p-adjusted value of less than 0.05. Volcano plots were generated using EnhancedVolcano v1.14.0^101^, with highlighted interferon stimulated genes (ISGs) being selected from the Molecular Signatures Database HALLMARK_INTERFERON_ALPHA_RESPONSE gene list^47^. Genes within this pathway were parsed from each virus infection and the stat values calculated using DESeq2 were used to compare the magnitude of change for each. A heatmap was generated using these values with Morpheus (https://software.broadinstitute.org/morpheus), and genes were organized using hierarchical clustering.

### Reverse transcriptase (RT)-quantitative PCR for validation of RNA sequencing data

Cells were lysed at indicated time points with buffer RLT Plus (Qiagen) and RNA was extracted according to manufacturer’s protocol. RNA was then reverse transcribed to produce cDNA using a High-Capacity cDNA Reverse Transcriptase Kit (Applied Biosystems, ThermoFisher). This cDNA was amplified using specific qRT-PCR primers for each target gene (primer sequences in **Table 2**). Fold changes in mRNA levels for indicated IFNs and ISGs were reported as fold changes over mock-treated cultures, using the formula 2^-Δ(ΔCt)^.

**Table 2.**
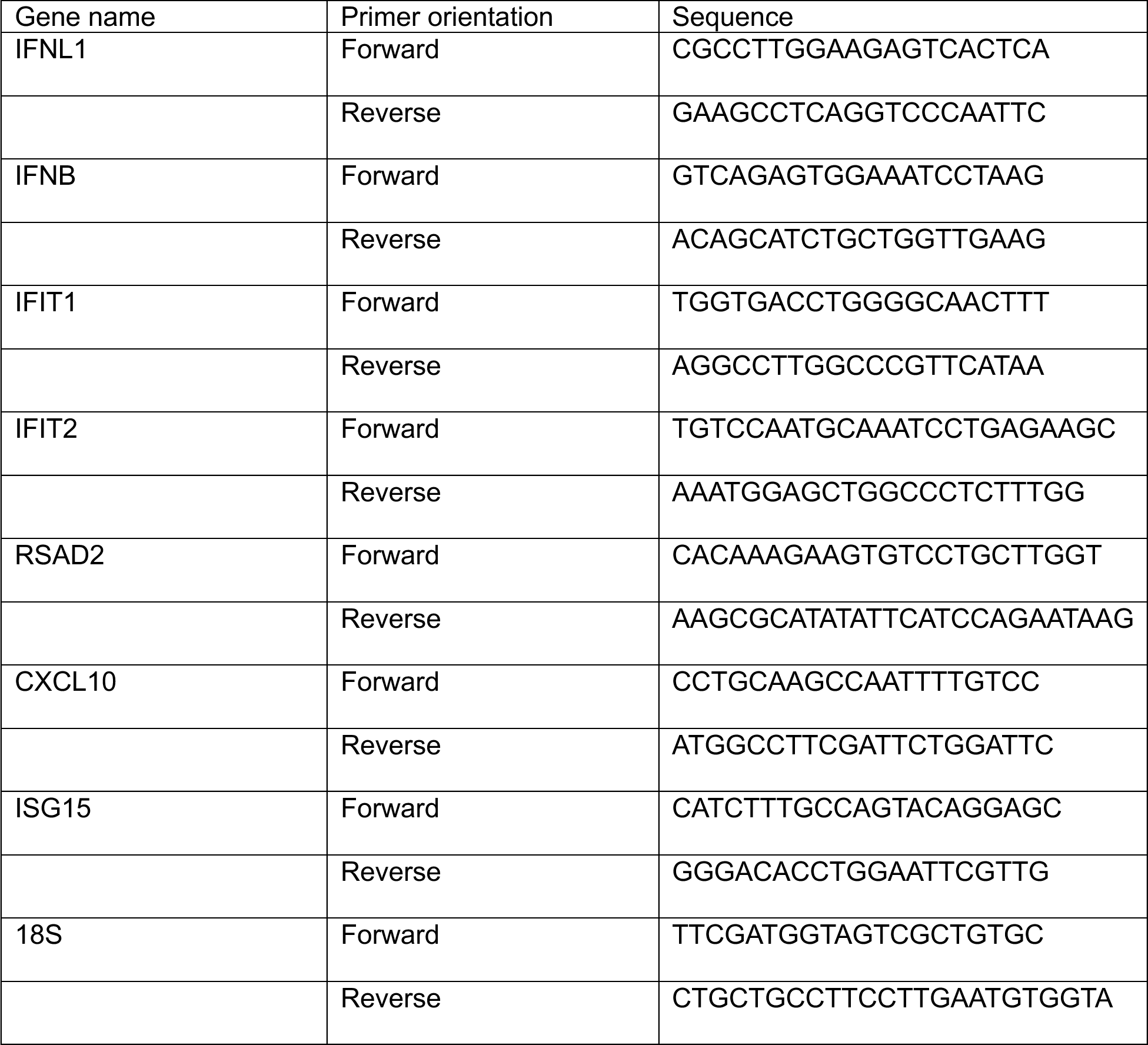
Primers used for qPCR analysis.

### Ruxolitinib treatments

Basal media of nasal ALI cultures was supplemented with 10 μM ruxolitinib (RUX) or DMSO (vehicle control). Basal media containing fresh RUX or DMSO was replaced at 0, 48, and 96 hpi. No detectable cytotoxicity was observed following RUX treatment as measured via lactate dehydrogenase assay.

### Measurement of trans-epithelial electrical resistance (TEER)

TEER was quantified using an EVOM ohm-voltmeter (World Precision Instruments) as previously described^40,95^. Briefly: TEER was quantified prior to infection and then measured after infection by placing each infected transwell into the Endohm-6 measurement chamber with PBS containing calcium and magnesium in both the apical and basal compartment. TEER measurements were reported as Ohms-cm^2 using the surface area of transwells (0.33cm^2^).

### Interferon pre-treatments

Nasal cultures were treated with 100 units/mL recombinant human IFN-β (Peprotech) or IFN-λ1 (BioLegend) 16 hours prior to infection (basally). Basal media was not changed following infection in IFN-treated cultures.

### Western blot analysis

Cell lysates were harvested at indicated time points using RIPA buffer (50mM Tris pH 8, 150mM NaCl, 0.5% deoxycholate, 0.1% SDS, 1% NP40) supplemented with protease inhibitors (Roche) and phosphatase inhibitors (Roche). Lysates were collected via scraping of entire surface of each transwell insert. Lysates were mixed 3:1 with 4X Laemmli sample buffer, boiled at 95°C for 10 minutes, and then separated via sodium dodecyl sulfate-polyacrylamide gel electrophoresis (SDS/PAGE) and transferred to polyvinylidene difluoride membranes. Blots were blocked in either 5% bovine serum albumin (BSA) or 5% nonfat milk and probed with antibodies as indicated in **Table 3**. Blots were stripped sequentially using Thermo Scientific Restore Western Blot stripping buffer.

**Table 3.** Antibodies used for western blotting.

| **Primary Antibody** | **Antibody species** | **Blocking buffer** | **Dilution** | **Catalog number** |
| --- | --- | --- | --- | --- |
| p-STAT-1 | Rabbit | 5% BSA in TBST | 1:1000 | Cell Signaling 7649 |
| STAT1 | Rabbit | 5% BSA in TBST | 1:1000 | Cell Signaling 9172 |
| IRF3 | Rabbit | 5% milk in TBST | 1:1000 | Cell Signaling 4302 |
| IFIT1 | Rabbit | 5% milk in TBST | 1:1000 | Cell Signaling 14769 |
| Viperin | Rabbit | 5% milk TBST | 1:1000 | Cell Signaling 13996 |
| MDA5 | Rabbit | 5% milk TBST | 1:1000 | Cell Signaling 5321 |
| p-PKR | Rabbit | 5% BSA in TBST | 1:1000 | Abcam 32036 |
| PKR | Rabbit | 5% milk TBST | 1:1000 | Cell Signaling 12297 |
| SARS-CoV-2 Nucleocapsid | Rabbit | 5% milk TBST | 1:2000 | Genetex GTX135357 |
| MERS-CoV Nucleocapsid | Mouse | 5% milk TBST | 1:2000 | Sino Biological 40068-MM10 |
| HCoV-229E Nucleocapsid | Rabbit | 5% milk TBST | 1:2000 | Sino Biological 40640-V07E |
| HCoV-NL63 Nucleocapsid | Rabbit | 5% milk TBST | 1:2000 | Sino Biological 40641-T62 |
| HRV-16 VP0/2 | Mouse | 5% milk TBST | 1:1000 | QED Biosciences 18758 |
| IAV HA | Mouse | 5% milk TBST | 1:2000 | Sigma H9658 |
| GAPDH | Rabbit | 5% milk or 5% BSA in TBST | 1:2000 | Cell Signaling 2118 |
| **Secondary Antibody** | **Antibody species** | **Blocking buffer** | **Dilution** | **Catalog number** |
| Anti-rabbit IgG HRP-linked | Goat | Same as primary | 1:3000 | Cell Signaling 7074 |
| Anti-mouse IgG HRP-linked | Horse | Same as primary | 1:3000 | Cell Signaling 7076 |

### Data Availability

Raw and processed RNA-seq data for all infection conditions will be deposited in the Gene Expression Omnibus database prior to publication. All other data are available upon request. Any material or related protocols mentioned in this work can additionally be obtained by contacting the corresponding author.

## Supplemental Figure Legends

**S1 RT-qPCR validation of bulk RNA-Seq results.** Nasal ALI cultures were infected with each virus (MOI = 1, 33°C), lysed at 96 hpi (SARS-CoV-2, MERS-CoV, HCoV-229E, HCoV-NL63) or 48 hpi (HRV-16, IAV), RNA extracted and analyzed by RT-qPCR with primers specific for indicated genes. Data are reported as fold changes relative to mock-infected cultures (calculated as 2^-Δ(ΔCt)^). Data shown are from one experiment representative of two experiments conducted in separate batches of pooled-donor nasal cultures.

**S2 Viral protein controls confirming infection by indicated viruses in Figure 2C**. Nasal ALI cultures were infected with each virus (MOI = 1, 33°C), lysed at 96 hpi (SARS-CoV-2, MERS-CoV, HCoV-229E, HCoV-NL63 N protein) or 48 hpi (HRV-16, IAV), and protein samples separated via SDS-PAGE followed by transfer on to a PVDF membrane for detection using indicated antibodies specific to surface proteins of each virus. This western blot corresponds to western blot shown in **Figure 2D**.

**S3 Temperature does not significantly impact replication of MERS-CoV or IAV.** Nasal ALI cultures were equilibrated at indicated temperature (33°C or 37°C) for 48 hours prior to infection by MERS-CoV (A) or IAV (B). ASL was collected at the indicated time points post infection and quantified via plaque assay. Averaged replication kinetics from infections in two independent sets of pooled-donor ALI cultures are shown in mean ± SD. Statistical significance of differences in titer at 33°C vs. 37°C for each virus was calculated by repeated measures two-way ANOVA and shown as a table: *, *P* ≤ 0.05. Comparisons that did not reach significance are not labeled.

**S4 Temperature-dependent IFN responses during HCoV-229E and HRV-16 infection.** Nasal ALI cultures were equilibrated at indicated temperature (33°C or 37°C) for 48 hours prior to infection by HCoV-229E (A) or HRV-16 (B). Western blot analysis was performed using lysates of cells harvested at indicated time points. Immunoblots were probed with antibodies against indicated proteins involved in the IFN signaling response. Data shown are from one representative of three independent experiments conducted in separate batches of pooled-donor nasal cultures. (C, D) Cultures were pre-treated with either RUX or DMSO at 10 µM in the basal media at the start of temperature equilibration 48 hours pre infection. Cultures were then infected with HCoV-229E (C) or HRV-16 (D) in triplicate (MOI = 1), ASL collected at indicated time points and infectious virus quantified by plaque assay. Average viral titer for is shown as mean ± SD. Statistical significance of differences in titer between each condition was calculated by repeated measures two-way ANOVA and shown as a table: *, *P* ≤ 0.05; **, *P* ≤ 0.01; ***, *P* ≤ 0.001; ****, *P* ≤ 0.0001. Data shown are from one representative of three independent experiments, each performed in triplicate using independent batches of pooled-donor cultures.

