## Supplement for "Interferon signaling in the nasal epithelium distinguishes among lethal and common cold respiratory viruses and is critical for viral clearance"

# S1

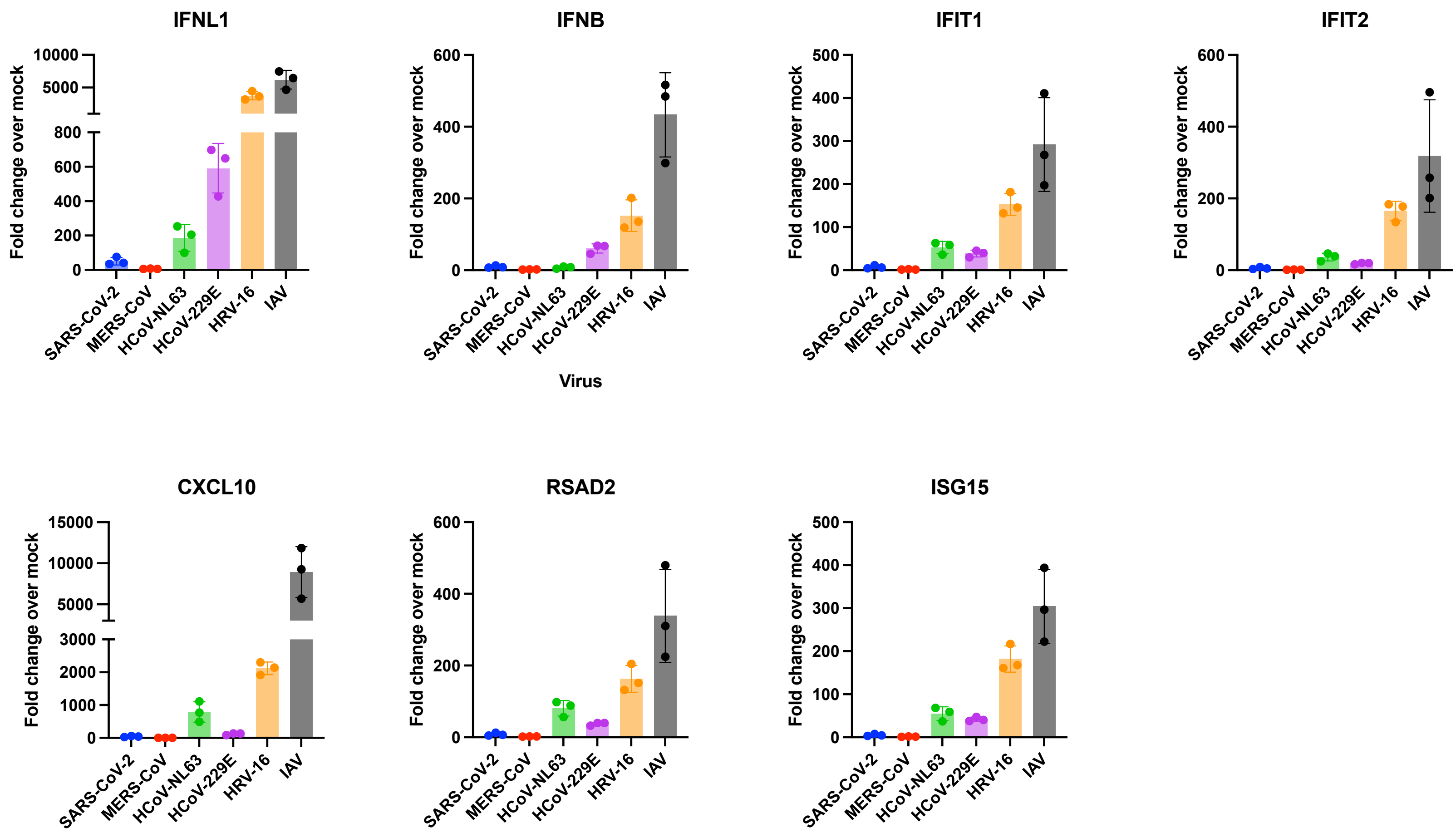
IFNL1
IFNB
IFIT1
IFIT2
CXCL10
RSAD2
ISG15
Fold change over mock
Virus
SARS-CoV-2
MERS-CoV
HCoV-NL63
HCoV-229E
HRV-16
IAV

**S2**

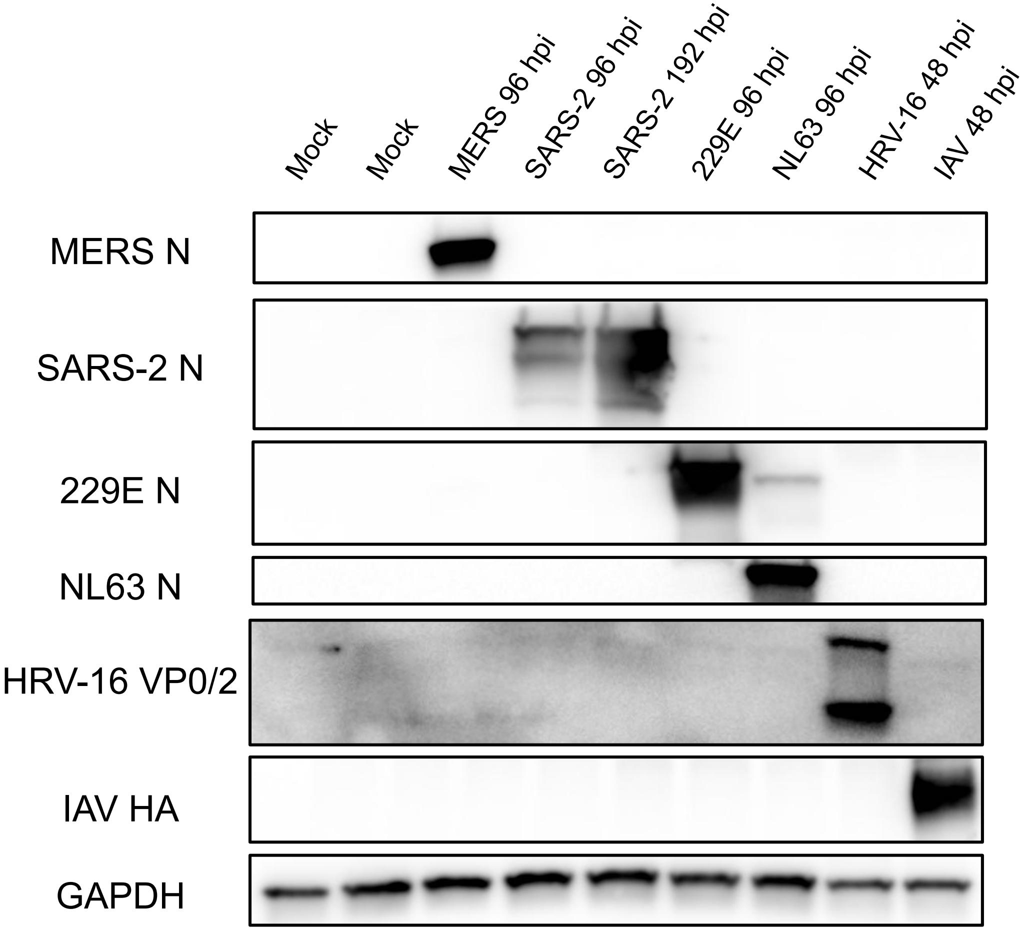
Mock
Mock
MERS 96 hpi
SARS-2 96 hpi
SARS-2 192 hpi
229E 96 hpi
NL63 96 hpi
HRV-16 48 hpi
IAV 48 hpi
MERS N
SARS-2 N
229E N
NL63 N
HRV-16 VP0/2
IAV HA
GAPDH

S3

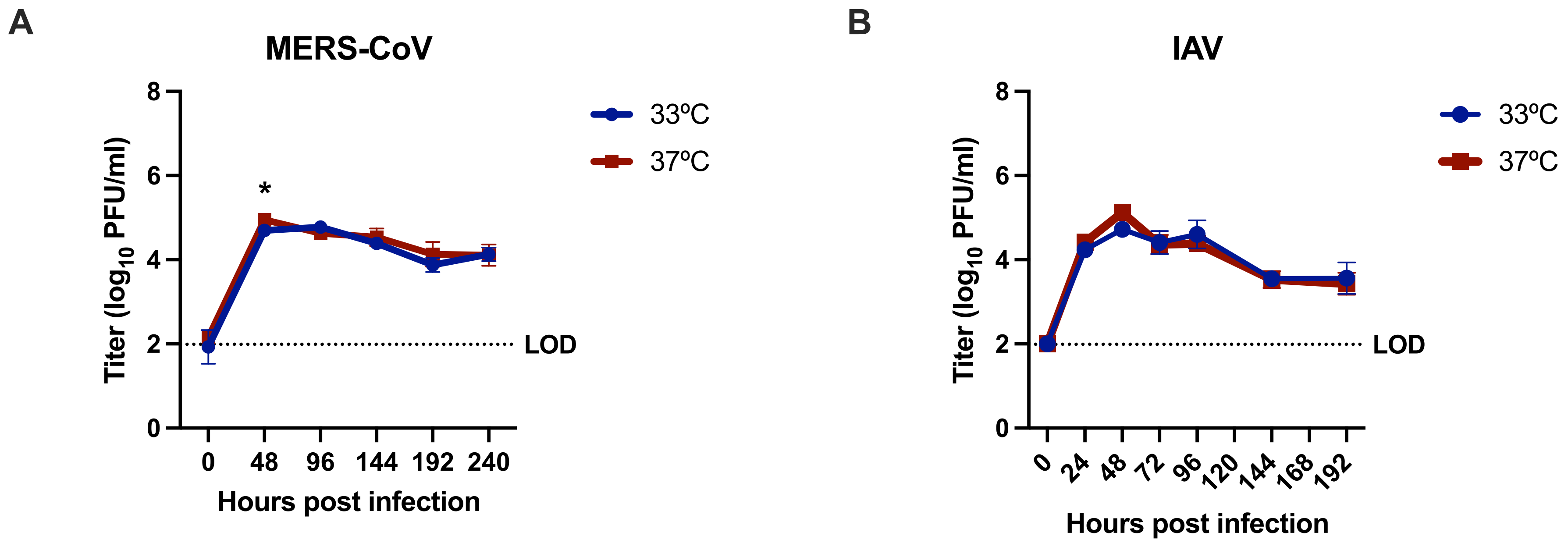
A
MERS-CoV
Titer (log10 PFU/ml)
8
6
4
2
0
*
LOD
0
48
96
144
192
240
Hours post infection
33ºC
37ºC
B
IAV
Titer (log10 PFU/ml)
8
6
4
2
0
LOD
0
24
48
72
96
120
144
168
192
Hours post infection
33ºC
37ºC

**S4**

**A**

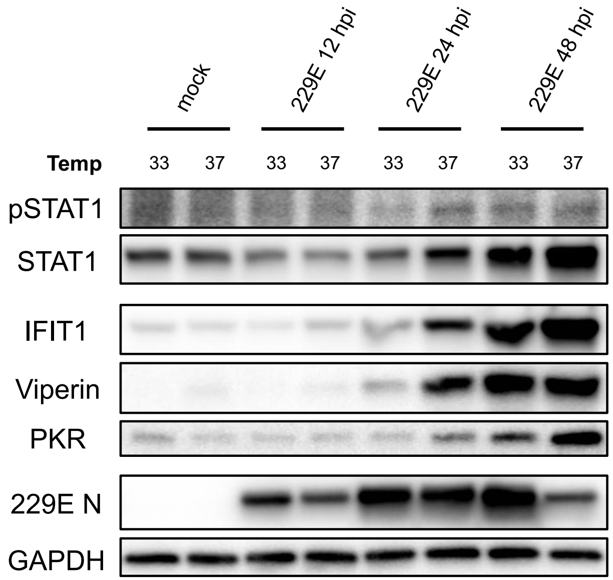


**C**

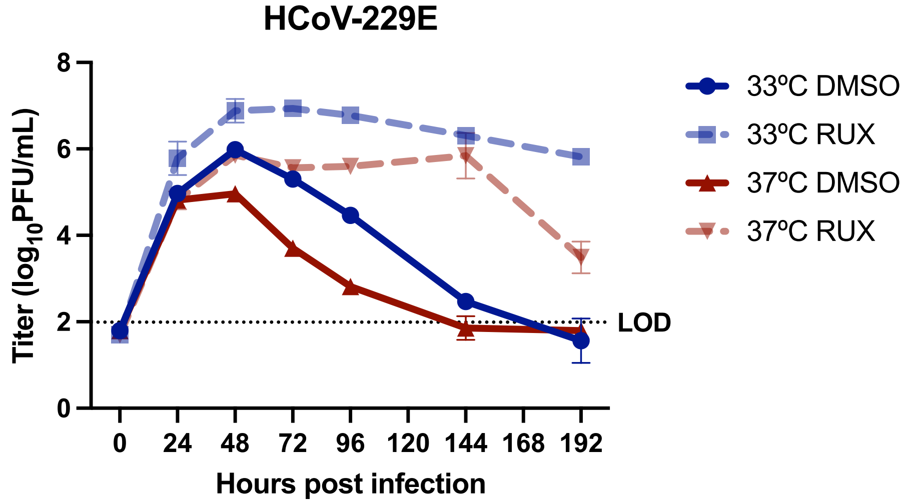


| Comparison | 0hpi | 24hpi | 48hpi | 72hpi | 96hpi | 144hpi | 192hpi |
|---|---|---|---|---|---|---|---|
| 33ºC DMSO vs. 33ºC RUX | ns | ns | * | **** | **** | **** | ** |
| 33ºC DMSO vs. 37ºC DMSO | ns | ns | ** | ** | ** | ns | ns |
| 33ºC DMSO vs. 37ºC RUX | ns | ns | ns | * | ** | ** | ** |
| 33ºC RUX vs. 37ºC DMSO | ns | * | ** | **** | **** | ** | *** |
| 33ºC RUX vs. 37ºC RUX | ns | * | * | **** | ** | ns | ** |
| 37ºC DMSO vs. 37ºC RUX | ns | ns | ** | *** | **** | ** | ** |

**B**

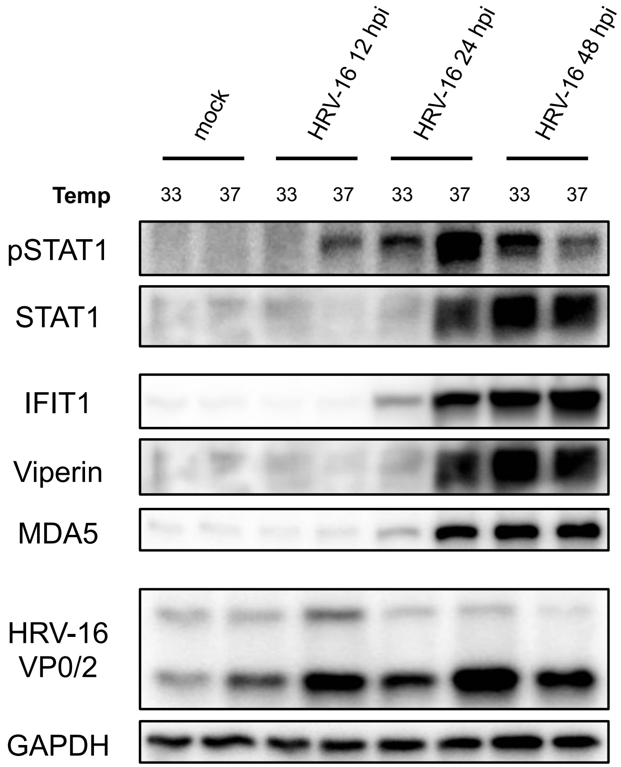


**D**

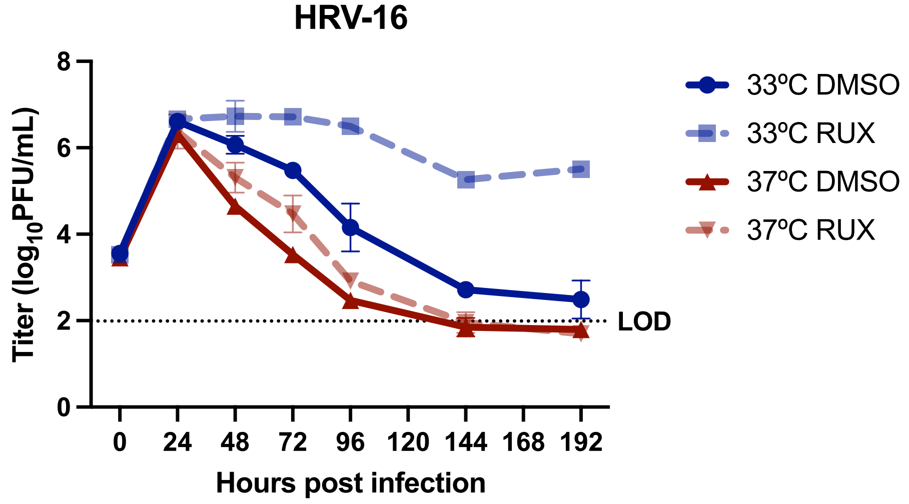


| Comparison | 0hpi | 24hpi | 48hpi | 72hpi | 96hpi | 144hpi | 192hpi |
|---|---|---|---|---|---|---|---|
| 33ºC DMSO vs. 33ºC RUX | ns | ns | ns | ** | * | **** | ** |
| 33ºC DMSO vs. 37ºC DMSO | ** | * | ** | ** | * | ns | ns |
| 33ºC DMSO vs. 37ºC RUX | * | ns | * | * | ns | * | ns |
| 33ºC RUX vs. 37ºC DMSO | * | ns | ** | **** | **** | ** | **** |
| 33ºC RUX vs. 37ºC RUX | ns | ns | ** | * | **** | *** | *** |
| 37ºC DMSO vs. 37ºC RUX | ns | ns | ns | ns | * | ns | ns |
